# Intracellular pH dynamics respond to extracellular matrix stiffening and mediate vasculogenic mimicry through β-catenin

**DOI:** 10.1101/2024.06.04.597454

**Authors:** Leah M. Lund, Angelina N. Marchi, Laura Alderfer, Eva Hall, Jacob Hammer, Riley Moremen, Ijeoma Asilebo, Keelan J. Trull, Donny Hanjaya-Putra, Katharine A. White

**Affiliations:** Department of Chemistry and Biochemistry, University of Notre Dame 251 Nieuwland Science Hall Notre Dame, IN 46556 USA; Harper Cancer Research Institute, University of Notre Dame 1234 N. Notre Dame Avenue South Bend, IN 46617 USA; Bioengineering Graduate Program, Aerospace and Mechanical Engineering, University of Notre Dame 2010G McCourtney Hall East Notre Dame, IN 46556 USA; Chemical and Biomolecular Engineering, University of Notre Dame 250 Nieuwland Hall Notre Dame, IN 46556 USA; Siemens Healthineers 430 S Beiger St. Mishawaka, IN 46544; Vivodyne Suite 775 601 Walnut Street Philadelphia PA 19106 USA; Lander University 231 Stanley Ave Greenwood, SC 29649

## Abstract

Dysregulated intracellular pH (pHi) dynamics and an altered tumor microenvironment have emerged as drivers of cancer cell phenotypes. However, the molecular integration between the physical properties of the microenvironment and dynamic intracellular signaling responses remains unclear. Here, we identify a mechanistic link between ECM stiffness and pHi dynamics in driving vasculogenic mimicry (VM), an aggressive cancer phenotype associated with poor prognosis. We performed single-cell imaging of pHi in lung and breast metastatic cell lines cultured on tunable-stiffness hydrogel systems. We used two tunable-stiffness hydrogel systems to independently model stiffness induced by increased protein secretion (Matrigel) and increased protein crosslinking (Hyaluronic acid gels). We show that increased ECM stiffness lowers single-cell pHi in both lung and breast metastatic cell lines. We also observed that stiff ECM promotes a distinct morphological phenotype called vasculogenic mimicry (VM). Importantly, we show that low pHi is a necessary mediator of VM, as raising pHi on stiff ECM reduces VM phenotypes. We also find that lowering pHi on soft ECM was sufficient to induce VM in the absence of extracellular stiffening. We characterized β-catenin as a pH-dependent molecular mediator of VM, where stiffness-driven increases in β-catenin abundance can be overridden by high pHi, which destabilizes β-catenin to reduce VM on stiff ECM. In contrast, the transcription factor FOXC2 is activated by ECM stiffness but is insensitive to pHi, and its activity alone is insufficient to maintain VM at high pHi when β-catenin is lost. We uncover a novel mechanotransduction axis in which ECM stiffness regulates intracellular pH to drive β-catenin-induced VM. We also show pHi dynamics can override mechanosensitive cell responses to the extracellular microenvironment. Thus, our work positions pHi as an integrator of mechanotransduction in cancer, suggesting a new framework for therapeutically targeting pHi in cancer and perhaps in other diseases driven by ECM remodeling.

## Introduction

The extracellular matrix (ECM) is a protein-rich structure that becomes dysregulated in cancer, driving cancer cell adaptation and the promotion of cancer cell phenotypes^1^. This increasingly rigid and dense tumor ECM promotes cancer cell invasion and vasculogenic mimicry, an adaptive cancer phenotype ^2,3^. In addition to the dysregulated extracellular environment, cancer cells also experience dysregulated pH dynamics^4^, with increased intracellular pH (pHi) (>7.4) and decreased extracellular pH (pHe) (<7.2) compared to normal epithelial cells (pHi 7.0-7.3; pHe 7.4)^5^. This reversal of the pH gradient is an early event in cellular transformation^6^ and has been directly linked to adaptive changes in cancer cell signaling, metabolism, proliferation, and evasion of apoptosis^5^.

Increased ECM stiffness promotes various cancer cell phenotypes including increased hypoxia^7^, vasculogenic mimicry^8^, cell durotaxis^9^, and selection for tumor-initiating cells or cancer stem-cell phenotypes^4,10–13^. Importantly, many equivalent or similar processes are also linked to dysregulated pHi dynamics, including hypoxia^11^, cell invasion^4^, and maintenance of a stem-like phenotype in adult and embryonic stem cell models^14^. However, the molecular mechanisms that integrate the physical properties of the microenvironment with intracellular cancer cell signaling are largely unknown.

While prior work has shown pHi dynamics can directly regulate normal mechanosensitive behaviors including focal adhesion remodeling^15^ and epithelial cell- cell contacts^16,17^, there are significant gaps in knowledge of the molecular crosstalk between ECM stiffening and pHi dynamics in cancer cells. One limitation is technical: it is challenging to develop mechanically tunable model systems that mimic physiological ECM dysregulation with suitable control. Previous studies have used synthetic ECM models, including Matrigel/Geltrex-^18^ and Hyaluronan-based^19^ systems. However, these studies have not decoupled the contributions of ECM protein abundance and ECM crosslinking density as independent drivers of mechanosensitive cell responses.

Another limiting factor in characterizing molecular links between ECM stiffness and pHi is that most mechanistic studies of pHi dynamics are performed under non-physiological culture conditions and lack single-cell resolution. These limitations are compounded when the goal is to explore effects of physical forces on cellular pHi dynamics and in the context of phenotypically heterogeneous cancer cells.

Here, we pair synthetic tunable-stiffness ECM models with live-cell pHi measurements and non-invasive pHi manipulation approaches to elucidate how pHi dynamics respond to ECM stiffening. We further explore the mechanistic role of pHi in regulating a cancer-associated mechanosensitive phenotype called vasculogenic mimicry (VM). We use two synthetic matrix models to mimic ECM stiffening through increasing protein abundance (Matrigel/Geltrex) and crosslinking density (hyaluronic acid gels), and measure single-cell pHi in metastatic breast and lung cancer cells. We show that pHi decreases with increased stiffness using both matrix models. We also demonstrate that cells plated on stiff ECM acquire distinct VM phenotypes that can be modulated by dynamically altering pHi. Importantly, raising pHi in cells plated on stiff matrix reduces VM phenotypes while lowering pHi in cells plated on soft matrix induces acquisition of a VM phenotype. We also investigate the pH dependence of two molecular regulators of VM (β-catenin and FOXC2). We show that β-catenin is a pH-dependent mediator of VM phenotype while FOXC2 activity is pH-insensitive. This suggests β-catenin as a necessary regulator of pH-dependent vasculogenic mimicry. Overall, our work reveals a previously unidentified link between mechanosensing and pHi dynamics in cancer and further suggests low pHi as a necessary and sufficient mediator of VM, a phenotype associated with aggressive cancers.

## Results

### Stiffening extracellular matrix lowers pHi in metastatic human lung carcinoma

Increased tumor microenvironment (TME) stiffness (>1000 Pa) can be caused by increased ECM protein deposition or by increased crosslinking of the ECM proteins compared to normal epithelial ECM (50-100 Pa)^20^ (**Figure 1A**). To investigate the central hypothesis of how a stiffening extracellular environment alters pHi, we used two tunable-stiffness hydrogel models to control ECM stiffness with high specificity. To mimic the effects of ECM stiffness changes resulting from altered ECM protein crosslinking, we used a hyaluronic-acid (HA) gel system where variable crosslinking density tunes ECM stiffness independently of protein concentration or composition^21,22^. HA is a non-sulfated linear polysaccharide of (1-β-4)d-glucuronic acid and (1-β-3)N-acetyl-d-glucosamine, and is a ubiquitous component of the ECM^19^. HA is abundant in the extracellular environment of the lung and brain^19^, and increased HA secretion is associated with cancer^19^ and fibrotic diseases of the liver and lung^23^.

**Figure 1:**
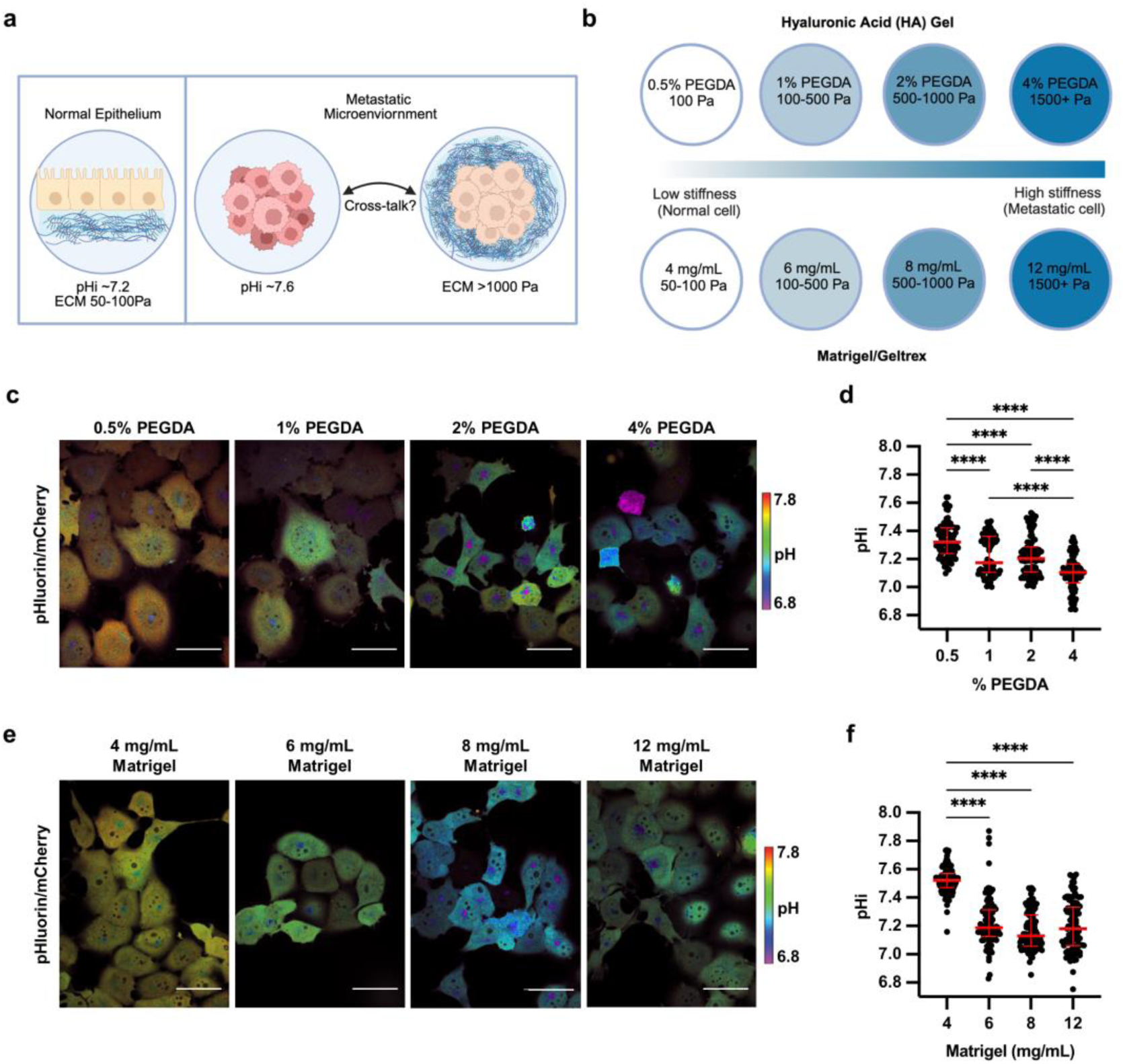
Stiffening extracellular matrix lowers pHi in metastatic human lung carcinoma (H1299). **a)** Schematic of increased pHi and ECM stiffening (via increased protein secretion and increased protein crosslinking) associated with tumorigenesis. **b)** Schematic of synthetic ECM models with tunable-stiffness (∼50 Pa-1,500 Pa). The Matrigel (or Geltrex) model mimics increased ECM protein secretion while hyaluronic acid (HA) gel system mimics increased ECM protein crosslinking. **c)** Representative images of H1299 cells stably expressing mCherry-pHluorin pH biosensor plated on varying HA gel stiffnesses. Images show ratiometric display of pHluorin/mCherry fluorescence. Scale bars: 50 μm. **d)** Quantification of single-cell pHi data collected as shown in (c). (n=3 biological replicates; n=91 0.5% PEGDA, n=90 1% PEGDA, n=102 2% PEGDA, n=89 4% PEGDA. Red lines show medians ± IQR). **e)** Representative images of H1299 cells stably expressing mCherry-pHluorin pH biosensor plated on varying Matrigel stiffnesses. Images show ratiometric display of pHluorin/mCherry fluorescence. Scale bars: 50 μm. **f)** Quantification of single-cell pHi data collected as shown in (e). (n=3 biological replicates; n=93 4mg/mL, n=92 6mg/mL, n=102 8mg/mL, n=97 12mg/mL. Red lines show medians ± IQR). For (d) and (f), significance was determined by a Kruskal-Wallis test (****P<0.0001).

Recent work has shown that HA can be functionalized to contain thiol-reactive cross-linkable regions, with high crosslinking adding rigidity to the ECM, enabling tunable stiffness^24^. Our HA gel tunable-stiffness model consists of a uniform concentration and mixture of gelatin and thiol-modified hyaluronan across stiffnesses, while stiffness is controlled by modulating amounts (%) of thiol-reactive PEGDA crosslinker (see methods for details). The HA gel model consists of four levels of crosslinking agent mimicking ECM stiffness changes induced by increased protein crosslinking and has a previously reported tunable stiffness range from ∼100-1500 Pa^25^ (**Figure 1B**). This system allows us to modulate matrix stiffness by adjusting the extent of ECM protein crosslinking while maintaining a consistent concentration of matrix components (hyaluronan and gelatin) across all stiffness conditions. This enables modeling ECM stiffness effects independently of matrix concentration changes.

To mimic the effects of stiffness changes induced by increased ECM protein secretion, we used Matrigel- or Geltrex-based tunable-stiffness gel systems. Matrigel and Geltrex are naturally-derived matrices that mimic the tumor microenvironment of stromal-rich tissues, such as breast, lung, and prostate^26^. The Matrigel and Geltrex commercial matrix mixtures are rich in laminin and collagen, ECM proteins that directly promote integrin signaling^27^. Varying the concentration of Matrigel and Geltrex effectively titrates ECM protein concentrations^28^, mimicking ECM stiffening caused by the increased secretion of ECM proteins^29^. We generated tunable-stiffness Matrigel/Geltrex models using four Matrigel/Geltrex concentrations (4 mg/mL-12 mg/mL) with stiffness ranges of ∼50-1,500 Pa^30–32^ (**Figure 1B**). For these stiffness determinations, the manufacturer reports a shear modulus (*G*′) that can be converted to Young’s modulus (matrix stiffness) using the following equation E= 2*G*’(1 + *v*). Prior work has indicated that hydrogels can be assumed to be incompressible^33,34^, such that their Poisson’s ratio (*v*) approaches 0.5, simplifying the equation to E=3*G*’. Importantly, in the Matrigel/Geltrex tunable-stiffness gel systems, as the ECM protein concentrations increase, so does the available ligand concentration for integrin-mediated interactions. This gel model allows the assessment of ECM stiffening on pHi when intracellular integrin signaling is also titrating.

With the two tunable-stiffness hydrogel systems established, we next selected cancer cell lines that originated from tissues with a relatively soft ECM, such as lung and breast, where tumorigenic ECM stiffening has been associated with both increased metastasis and invasion^26^. We have previously established and characterized single-cell pHi heterogeneity in a clonal metastatic lung cancer cell line (H1299) and a clonal breast cancer cell line (MDA-MB-231), all plated and imaged on glass^35^. We have engineered these cell lines to stably express a genetically-encoded ratiometric pH biosensor mCherry-pHluorin (mCh-pHl)^35^. This biosensor is a fusion of the fluorescent protein pHluorin (pKa 7.1) that is pH-sensitive in the physiological range, and the fluorescent protein mCherry, that is pH-insensitive in the physiological range^36^. For accurate pHi measurements in single cells, ratiometric imaging of pHluorin and mCherry fluorescence can be performed followed by single-cell standardization using isotonic buffers with a known pH containing the protonophore Nigericin to equilibrate intracellular and extracellular (buffer) pH^37^. Single-cell standard curves are then generated, enabling back-calculation of pHi from pHluorin and mCherry fluorescence intensity ratios (**Figure S1**, see methods for details). This biosensor has successfully been used in prior studies to measure single-cell spatiotemporal pHi dynamics in clonal cancer and normal epithelial cell populations without affecting cell morphology or behavior^15,35,36^.

To determine the effects of altered ECM stiffness on pHi, we cultured H1299 cells expressing the mCh-pHl biosensor on matrix-coated imaging dishes for 48 hours. This incubation allowed for cells to adhere and respond to the varied stiffness of each matrix system. In cells plated on HA gels, single-cell pHi decreased with increasing stiffness (**Figure 1C**). Cells plated on the stiffest matrix (4% PEGDA) had significantly decreased pHi (**Figure 1D**; 7.10±0.07; median±interquartile range (IQR)) compared to cells on the softest matrix (0.5% PEGDA) (**Figure 1D**; 7.32±0.10; median±IQR). We also observed that intermediate ECM stiffnesses (1% PEGDA and 2% PEGDA) produced intermediate pHi values, with a stepwise trend of decreasing pHi with increasing stiffness (**Figure 1D**; 2% PEGDA 7.20±0.10; 1% PEGDA 7.17±0.19; medians±IQR). The overall decrease in pHi of ∼0.2 pH units between soft and stiff ECM is within the range of physiological pHi dynamics shown to regulate normal cell behaviors, including cell cycle progression^35^, differentiation^13,38^, and migration^39^. This result demonstrates that ECM stiffening through changes in protein crosslinking drives significant decreases in single-cell pHi of clonal metastatic lung cancer cells. These data suggest that progressive changes in ECM stiffness within the physiological range of normal to metastatic mechanical stiffness environments can alter pHi in metastatic cancer cells, suggesting a potential role for pHi in mechanosensitive cancer cell signaling and behaviors.

We next determined whether the stiff ECM decreased pHi using the Matrigel tunable-stiffness models, where ECM protein concentration is the predominant driver of altered stiffness. In cells plated on varied Matrigel stiffnesses, single-cell pHi decreased with increasing stiffness (**Figure 1E**). Cells plated on the stiffest matrix (12 mg/mL) had a significantly decreased pHi (**Figure 1F**; 7.18±0.15; median±IQR) compared to the softest matrix (**Figure 1F**; 4 mg/mL; 7.52±0.49; median±IQR). The decrease in pHi of ∼0.35 units between stiffest (∼1,500 Pa) and softest (∼50 Pa) ECM in this system is also consistent with the pHi changes measured between stiffest and softest HA gel models (∼0.22 units). However, in the Matrigel tunable-stiffness model system, the pHi of cells measured on intermediate stiffnesses (**Figure 1F**; 6 mg/mL Matrigel, 7.19±0.13; 8 mg/mL Matrigel, 7.13±0.14; medians±IQR) was not significantly different from the pHi of cells plated on a stiff matrix, suggesting that in the context of titrating integrin signaling, the effect may be more binary compared to stiffness changes induced by crosslinking. These results show that ECM stiffening lowers pHi in metastatic cells regardless of whether the stiffness change is induced by increased ECM protein abundance or ECM protein crosslinking.

We next confirmed that increased ECM stiffness decreases pHi using another metastatic breast cell model (MDA-MB-231), which is also derived from a stromal-rich environment (like H1299). We found that the pHi of MDA-MB-231 cells was decreased by ∼0.2 units in cells plated on a stiff matrix compared to soft matrix in both the HA gel (soft 7.43±0.14; stiff 7.28±0.10; medians±IQR) (**Figure S2A, B**) and the Matrigel (soft 7.40±0.14; stiff 7.20±0.13; medians±IQR) models (**Figure S2 C,D**). Again, with the HA gel model, we observed step-wise effects of pHi in response to stiffness with intermediate pHi values on the intermediate stiffnesses (**Figure S2B**), just as we saw with the metastatic lung model (**Figure 1C,D**). While the pHi of MDA-MB-231 cells was significantly decreased on stiff (12 mg/mL) compared to the softest (4 mg/mL), the pHi response of MDA-MB-231 on the intermediate stiffnesses was more variable on Matrigel (**Figure S2D**). While we cannot rule out that pHi response to ECM may vary depending on cell type or matrix composition, we predict that these discrepancies are due instead to cell-line dependent responses to the very low integrin ligand concentration when using the lowest dilution (4 mg/mL) of Matrigel/Geltrex. When using the HA-gel system, where protein concentration is maintained across stiffnesses, the behavior of H1299 and MDA-MB-231 are in close concordance.

Both H1299 and MDA-MB-231 are derived from soft tissue (breast and lung) and were isolated from soft metastatic sites (lymph node and pleural effusion). To determine whether metastatic cells from a stiff tissue of origin are similarly sensitive to ECM stiffening, we cultured metastatic bone cancer cells (U-2 OS) on soft and stiff HA gels. We found that these cells similarly had reduced pHi when cultured on stiff ECM compared to soft (**Figure S3**, 0.5% PEGDA. 7.28±0.09; 4% PEGDA 7.15±0.05; medians±IQR). This matched relationship of pHi and ECM stiffness in metastatic cells derived from both soft and stiff tissues suggest that decreased pHi may be a conserved metastatic cell response to ECM stiffening.

Taken together, these data show that increased ECM stiffness mediated by either increased crosslinking (HA gel model) or by increased ECM protein secretion (Matrigel model) decreases pHi of metastatic cancer cells. Our data also show that the stiffness-dependent decreases in pHi are not tissue-specific, as both breast and lung metastatic models exhibited lower pHi on stiff compared to soft matrices. In summary, these data demonstrate an inverse relationship between ECM stiffening and pHi in metastatic cell models and suggest a role for pHi in cell responses to mechanical ECM cues.

### Stiffness-dependent vasculogenic mimicry is reduced in high pHi conditions in metastatic cell models

When performing single-cell pHi measurements, we also observed a distinct change in overall cancer cell morphology that correlated with increased ECM stiffness. Metastatic cancer cells plated on soft matrix grew in flat lawns of cobblestone (H1299) or spindle-shaped cells (MDA-MB-231), forming a near-confluent sheet. However, on stiff matrix, the metastatic cancer cells grew in compact clusters of irregularly shaped cells, frequently exhibited 3D growth phenotypes, and formed connected bridges of elongated spindle shaped cells between 3D “nodes” (**Figure S4**). Similar changes in cell morphology are found in vasculogenic mimicry (VM), an aggressive cancer phenotype observed both *in vivo* and *in vitro*, where tumor cell organize into vessel-like structures, allowing nutrients and oxygen access independent of traditional angiogenesis^40^. Previous studies have shown that increased ECM stiffness drives VM^41^ phenotypes and characterized 2D VM phenotypes as a growth pattern where cells form distinct networks of tightly packed cells with surrounding open space devoid of cell growth^42^.

Our data show stiff ECM lowers single-cell pHi and induces VM phenotypes, leading to the hypothesis that low pHi is a necessary mediator of VM (**Figure 2A**). If this hypothesis is correct, raising pHi in cells plated on stiff matrix should reduce the VM phenotype (**Figure 2A**). To directly test this hypothesis, we established protocols to experimentally raise pHi in cells plated on stiff ECM. Prior work showed that 50 mM sodium bicarbonate supplemented into the media for 24 hours was sufficient to raise pHi in H1299 cells plated on glass^35^. We imaged single-cell pHi in H1299 cells plated on soft ECM, stiff ECM, and stiff ECM with bicarbonate supplementation to raise pHi (**Figure 2B**). We found that bicarbonate significantly increased pHi of cells plated on stiff ECM compared to untreated cells on stiff matrix (stiff 7.27±0.08; stiff + Bicarbonate 7.43±0.08; medians±IQR) (**Figure 2C).** The bicarbonate treatment increased the pHi of cells plated on the stiffest ECM by approximately 0.2 pH units (**Figure 2C**), which is similar to the magnitude of pHi changes we observed between soft and stiff ECM across the various cell lines and gel systems.

**Figure 2:**
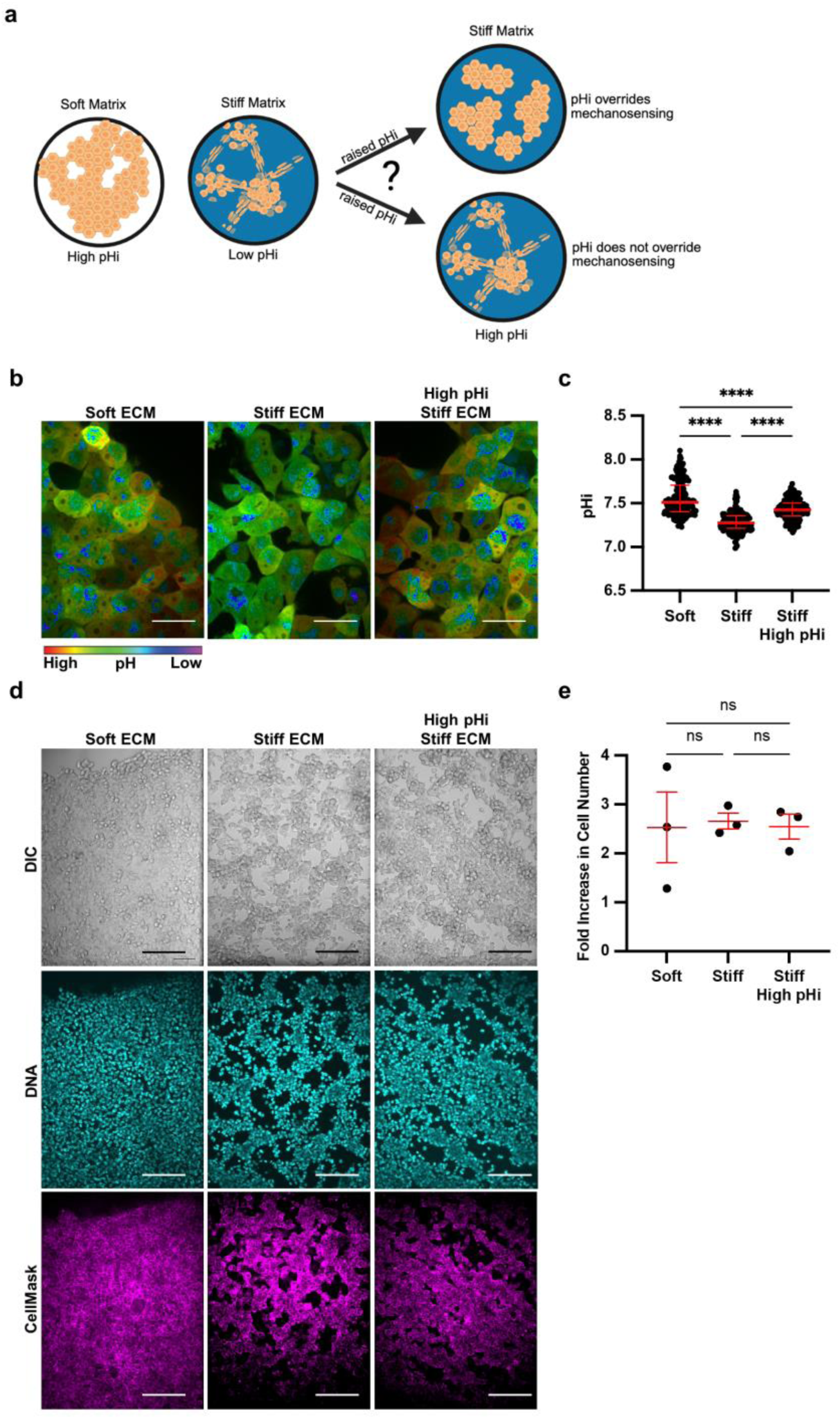
Stiffness-dependent vasculogenic mimicry is reduced when pHi is increased on stiff ECM. **a)** Schematic of vasculogenic mimicry (VM) in 2D on stiffening matrix. **b)** Representative images of H1299 cells stably expressing mCherry-pHluorin pH biosensor plated on soft (0.5% PEGDA) and stiff (4% PEGDA) HA gels and stiff (4% PEGDA) with raised pHi (culture media supplementation with 50mM sodium bicarbonate). Images show ratiometric display of pHluorin/mCherry fluorescence ratios. Scale bars: 50 μm. **c)** Quantification of single-cell pHi data collected as shown in (b) (n=3 biological replicates; n=201 soft, n=237 stiff, n=239 stiff high pHi. Red lines show medians ± IQR). **d)** Representative images of H1299 cells plated on soft (0.5% PEGDA) and stiff (4% PEGDA) HA gels. Images show differential interference contrast (DIC) and Hoechst stain (DNA, cyan). Scale bars: 100 μm. **e)** Quantification of cell proliferation across manipulation conditions. (n=3 biological replicates. Red lines show means ± SEM).

We next tested the effects of increased pHi on the stiffness-dependent vasculogenic mimicry phenotype. We found that H1299 cells acquired a vasculogenic mimicry phenotype on stiff matrix, and this VM phenotype was abrogated when pHi was increased (**Figure 2D**). Cells plated on stiff matrix with bicarbonate-induced increases in pHi grew in a 2D cobblestone-like morphology similar to that of cells grown on soft ECM (**Figure 2D**, additional representative images in **Figure S4**). To confirm that the observed pH-dependent changes in cell morphology were not due to differences in cell proliferation, we assayed proliferation rates in H1299 cells plated on soft and stiff ECM with and without increased pHi. Importantly, we did not observe any significant differences in proliferation rates across our experimental conditions over the timeframe tested (**Figure 2E**).

To quantify the observed stiffness- and pHi-dependent changes in cell morphology, we used a cell membrane marker and quantitative image analysis pipeline (see methods for details) to assess cell area (**Figure 3A**). Notably, single-cell area of H1299 cells was significantly lower in cells plated on stiff ECM compared to soft ECM (Figure 3B; stiff 339 µm^2^±189; soft 407 µm^2^±225; medians±IQR). This result demonstrates that cell area is a robust quantitative indicator that decreases with the acquisition of VM morphology phenotype on stiff ECM. This allows us to quantitatively distinguish cell morphologies corresponding to low VM and high VM using cell area. Importantly, we found that cell area significantly increased (**Figure 3B**; 374 µm^2^±193; median±IQR) when pHi was raised in H1299 cells plated on stiff ECM compared to control H1299 on stiff ECM (**Figure 3B**). This indicates that increased pHi significantly reduces the stiffness-dependent VM phenotype. The loss of VM networks and increased cell area when pHi is raised on stiff ECM demonstrates that low pHi is required for cells to maintain VM on stiff ECM.

**Figure 3:**
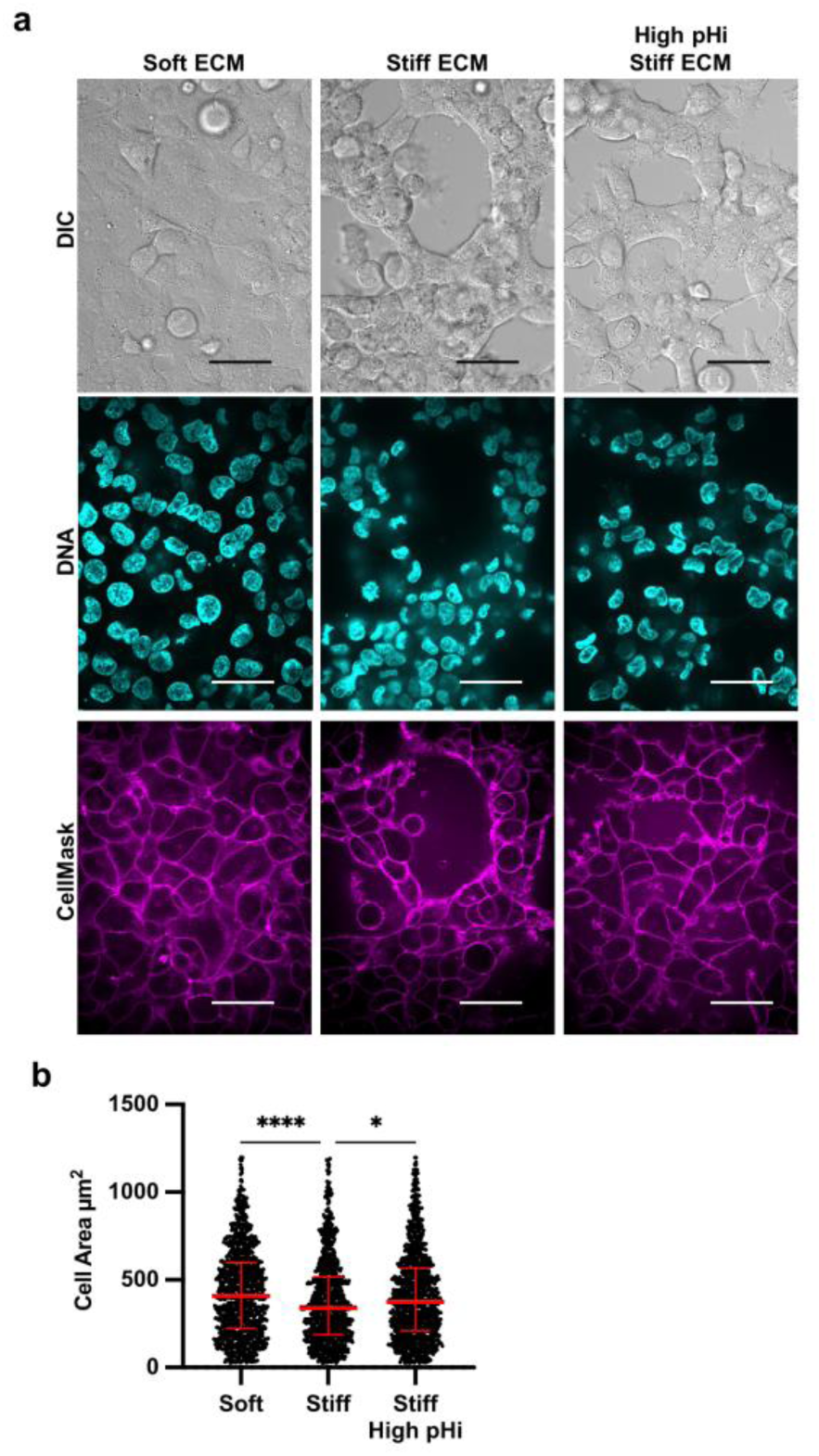
Vasculogenic mimicry phenotype decreases cell area on stiff ECM, which is rescued by increasing pHi in metastatic lung carcinoma. **a)** Representative images of H1299 cells plated on soft (0.5% PEGDA) and stiff (4% PEGDA) HA gels and stiff (4% PEGDA) with raised pHi. Images show differential interference contrast (DIC), Hoechst 33342 (DNA, cyan) and CellMask Deep Red membrane stain (Cy5, magenta). Scale bars: 50 μm. **b)** Quantification of single-cell area collected as shown in (a) (n=3 biological replicates, n=1061 soft, n=954 stiff, n=1078 stiff high pHi. Red lines show medians ± IQR).

We also probed the VM phenotype in MDA-MB-231 cells and U-2 OS cells and found that single-cell area of both MDA-MB-231 (**Figure S5**) and U-2 OS cells (**Figure S6**) was decreased in cells plated on stiff ECM compared to soft. Raising pHi in MDA-MB-231 cells on stiff ECM did produce a noticeable shift in morphology, reverting cells to the flatter, spread morphology observed on soft ECM (**Figure S5C**) and when we quantified cell area we found a trend toward increased cell size, though it was not statistically significantly different from stiff (**Figure S5D**, p=0.2549). However, MDA-MB- 231 cells already have pronounced mesenchymal-like phenotype which may obscure these cell area calculations (reducing the measurable dynamic range). These findings demonstrate that while high pHi can reverse stiffness-induced VM in metastatic cells, inherent mesenchymal traits of some metastatic cells like MDA-MB-231 may limit the extent to which VM-associated morphological change can be measured. Together, our findings confirm that vasculogenic mimicry is an ECM stiffness-mediated phenotype and further identify decreased pHi as a previously unrecognized necessary regulator of VM phenotypes.

### β-catenin abundance is stiffness-dependent, pHi-dependent, and necessary for stiffness-dependent vasculogenic mimicry

We next investigated molecular drivers of pH-dependent regulation of VM. In epithelial cells, VM is regulated by several mechanisms, including the activity and abundance of β-catenin^43,44^, a multifunctional protein that regulates cell-cell adhesion and transcription, and the activity of transcription factor FOXC2^42,45^. These two molecular regulators of VM have previously characterized pH-sensitive activity in epithelial models (β-catenin)^17,46^ or *in vitro* (FOXC2)^47^. However, existing literature is conflicting as to whether FOXC2 and β-catenin are truly independent drivers of VM. For example, prior data suggests that β-catenin functions upstream of FOXC2 in VM, with β-catenin directly controlling expression of FOX transcription factors^48^. However, other data suggests that FOXC2 can itself modulate Wnt signaling^49,50^ and rescue acquisition of vasculogenic mimicry when β-catenin levels are reduced^48^. We next used our models to differentiate roles of FOXC2 and β-catenin in regulating stiffness- and pH-dependent VM phenotypes.

While FOXC2 has been shown to be required for VM^45^, and sufficient to drive endothelial cell vascularization^48^, it is unclear whether FOXC2 abundance or activity is a sufficient driver of VM phenotypes. We first measured FOXC2 abundance in H1299 cells plated on soft ECM and stiff ECM and found no difference in FOXC2 expression (**Figure S7A,B**), indicating FOXC2 protein abundance is not regulated by stiffness or pHi. We next measured FOXC2 transcriptional activity in single cells using a FOXC2 transcriptional activity reporter plasmid (FOXC2-TAG-Puro) that has FOXC2 specific tandem repeats flanking a core DNA binding element upstream of GFP (LipExoGen, see methods) (**Figure S7C**). We found that FOXC2 activity was significantly increased in cells plated on stiff ECM compared to cells plated on soft ECM (**Figure S7D)**, suggesting that ECM stiffening is sufficient to increase FOXC2 transcriptional activity. However, FOXC2 transcriptional activity remained high when pHi was increased on a stiff matrix (**Figure S7**), suggesting pHi is not a sufficient modulator of FOXC2 activity in this model. Furthermore, our data show that high FOXC2 transcriptional activation is not a sufficient driver of VM in this model, as high pHi abrogates VM phenotypes (**Figure 2&3**) even as FOXC2 activity remains high (**Figure S7**).

We next tested the hypothesis that stiffness-associated pHi dynamics modulate VM by regulating β-catenin abundance in metastatic cancer cell lines. Previous work has shown that β-catenin abundance and transcriptional activity directly regulate VM^43^. Furthermore, increased β-catenin nuclear localization correlates with VM formation in colon cancer models^44^ and is associated with ECM stiffening in liver cancer models^51^. Importantly, ECM stiffening has also been shown to increase whole-cell β-catenin abundance in some cell lines, including human mesenchymal stem cells^52,53^. Work from our lab has also shown that pHi can directly regulate β-catenin stability in normal epithelial cells, with high pHi reducing β-catenin stability while low pHi stabilizes β- catenin and increases its adhesion and transcriptional activity^17^. However, our prior work did not assess pH-dependent β-catenin stability in non-epithelial models and did not fully characterize the functional consequences of pH-dependent β-catenin stability on cell behaviors.

While previous studies have demonstrated the role of β-catenin in regulating VM, these studies have not characterized the cellular cues by which a stiff ECM increases β- catenin abundance or nuclear localization. To determine the effect of ECM stiffening on β-catenin abundance, we performed immunofluorescent staining of β-catenin in H1299 cells plated on soft ECM and stiff ECM with and without pHi manipulation (**Figure 4A**). We found that both whole-cell and nuclear β-catenin abundance were significantly increased in cells plated on stiff ECM (low pHi) compared to cells plated on soft ECM (high pHi) (**Figure 4B,C**). This is in agreement with our prior work showing that low pHi stabilizes β-catenin in epithelial cells^53^. Furthermore, we found that when we raised pHi in cells plated on stiff ECM, VM was reduced, and whole-cell and nuclear β-catenin abundance was significantly reduced compared to control cells plated on stiff ECM (**Figure 4B,C**). We also confirmed these results in the MDA-MB-231 model, showing that low pHi on stiff ECM increases β-catenin whole cell and nuclear abundance compared to soft, and that raising pHi on stiff ECM reduces β-catenin abundance (**Figure S8**). Thus, our data demonstrate that increased β-catenin abundance is correlated with low pHi in human metastatic cancer cells. Taken together, these data suggest that pH-dependent β-catenin abundance underlies the observed pH-dependent regulation of VM. Finally, these data show that high pHi can override mechanosensing by decreasing β-catenin abundance, suggesting that low pHi functions as a necessary mediator of VM in cancer cells via stabilization of β-catenin.

**Figure 4:**
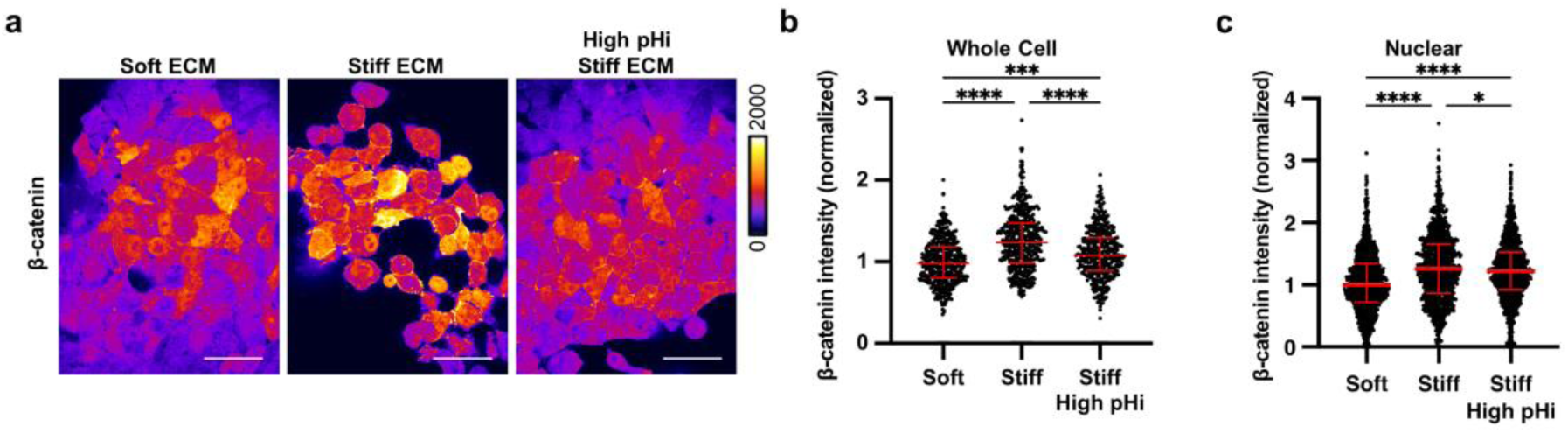
Increased pHi reduces β-catenin abundance and nuclear localization in stiff matrix conditions. **a)** Representative images of H1299 cells plated on soft (0.5% PEGDA), stiff (4% PEGDA) and stiff with raised pHi (4% PEGDA) HA gels fixed and stained for β-catenin. β-catenin is pseudocolored according to scale. Scale bars: 50 μm. **b)** Quantification of whole cell β-catenin intensity collected as shown in (a). (n=3 biological replicates, n=452 soft, n=486 stiff, n=415 stiff high pHi. Red lines show medians ± IQR). **c)** Quantification of nuclear β-catenin intensity collected as described in (a). (n=3 biological replicates, n=1043 soft, n=975 stiff, n=1157 stiff high. Red lines show medians ± IQR). For (b) and (c), significance was determined by a Kruskal-Wallis test (\**P*<0.05; \*\**P*<0.01; \*\*\**P*<0. 001; \*\*\*\**P*<0.0001).

We next asked whether stabilizing β-catenin at high pHi could rescue VM on stiff ECM. To test this, we stabilized β-catenin abundance in high pHi conditions using CHIR, an inhibitor of GSK-3β, a required component of the β-catenin destruction complex^54^. We confirmed that CHIR treatment does not disrupt the previously characterized stiffness-dependent and treatment-dependent pHi dynamics in H1299 lung cancer models (**Figure 5A**). Again, we observed reduced β-catenin on stiff ECM when pHi was raised, but CHIR supplementation to this condition significantly increased β-catenin abundance (**Figure 5B,C**). Furthermore, we found that when treated with the CHIR inhibitor, cells with raised pHi on a stiff matrix exhibited an increase in vascular mimicry. (**Figure 5D,E**). This data suggests that rescuing β-catenin abundance under high pHi conditions rescues VM on stiff ECM, and further demonstrates β-catenin as a primary regulator of pH-dependent VM in this model. Combined with our FOXC2 results, our data also show that β-catenin loss at high pHi regulates VM independently of FOXC2 activity, reducing VM phenotypes even while FOXC2 transcriptional activity remains high.

**Figure 5:**
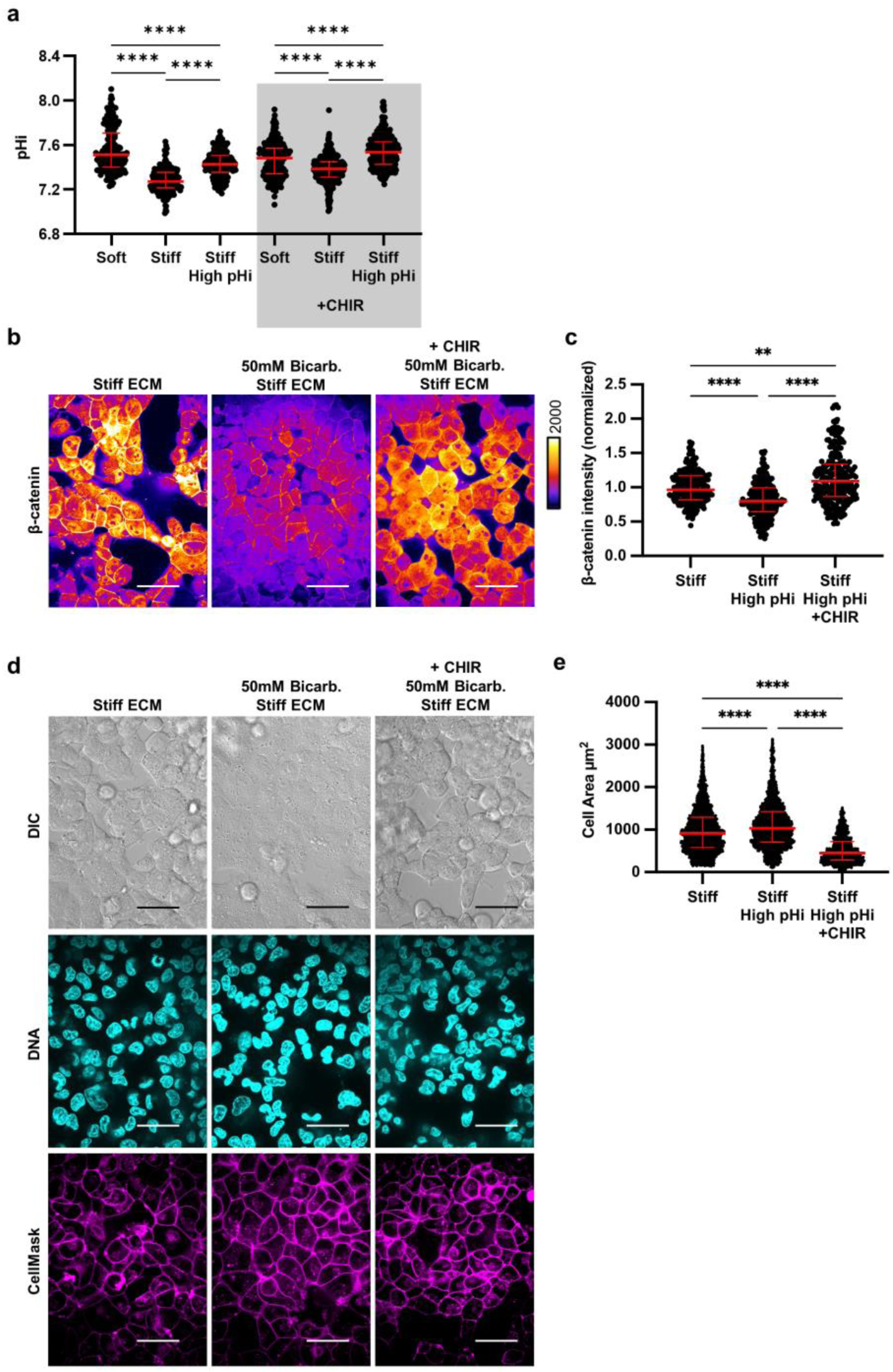
Stabilization of β-catenin abundance under high pHi rescues VM on a stiff matrix. **a)** Quantification of pHi data from parental H1299 cells and H1299 cells treated with CHIR treatment with and without treatment with sodium bicarbonate (Bicarb.) (see methods) (n=3 biological replicates. n=191 soft, n=219 stiff, n=198 stiff high pHi, n=302 soft +CHIR, n=373 stiff +CHIR, n=408 stiff high pHi +CHIR. Red lines show means ± SEM). **b)** Representative images of H1299 cells plated on stiff HA gels (4% PEGDA) with and without raised pHi (Bicarb.) and with or without CHIR treatment fixed and stained for β-catenin. β-catenin is pseudocolored according to scale. Scale bars: 50 μm. **c**) Quantification of whole cell β-catenin intensity collected as shown in (b). (n=3 biological replicates, n=452 stiff, n=486 stiff high pHi, n=415 stiff high pHi +CHIR. Red lines show medians ± IQR). **d)** Representative images of H1299 cells plated on stiff HA gels (4% PEGDA) with and without raised pHi (Bicarb.) and with or without CHIR treatment. Images show differential interference contrast (DIC), Hoechst 33342 (DNA, cyan) and CellMask Deep Red membrane stain (Cy5, magenta). Scale bars: 50 μm. **e)** Quantification of single-cell area collected as shown in (d) (n=3 biological replicates, n=452 stiff, n=486 stiff high pHi, n=415 stiff high pHi +CHIR. Red lines show medians ± IQR).

### Low pHi is a sufficient driver of vasculogenic mimicry phenotypes on soft ECM

The prior results suggest that low pHi is necessary for VM on stiff ECM, as raising pHi can override stiffness-associated VM phenotypes. We next hypothesized that low pHi is a sufficient mediator of VM and that lowering pHi in H1299 cells plated on soft matrix can induce stiffness-independent acquisition of VM phenotypes. To directly test this hypothesis, we used an H1299 cell line deficient in the sodium proton exchanger (H1299-NHE1 K.O., see methods). We performed an acid load recovery assay to confirm that the H1299-NHE1 K.O. cell line had no measurable NHE1 activity (**Figure S9**). This H1299-NHE1 K.O. cell line has significantly decreased basal pHi compared to parental H1299, and bicarbonate treatment raised pHi in this model (**Figure 6A**). Using this experimental system, we tested the effects of decreased pHi on modulating VM phenotypes. We found that the H1299-NHE1 K.O. cells acquired a VM phenotype on soft matrix, suggesting low pHi is indeed a sufficient driver of VM in the absence of stiff ECM mechanical cues (**Figure 6B**). Importantly, the stiffness-independent VM phenotype observed in H1299-NHE1 K.O. cells on soft ECM was again reduced when pHi was raised in cells on the soft matrix (**Figure 6B**). We used cell area to quantify the extent of VM phenotype and found that when pHi is lowered (H1299-NHE1 K.O.) in cells plated on soft ECM, single-cell area is significantly decreased compared to when pHi was raised in the same cells plated on soft ECM (**Figure 6C,D**). Our findings demonstrate that decreased pHi is sufficient to drive a VM phenotype in the absence of stiffening ECM mechanical cues. We confirm in this model that increasing pHi is sufficient to override VM phenotypes, even when VM is aberrantly generated on soft ECM.

**Figure 6:**
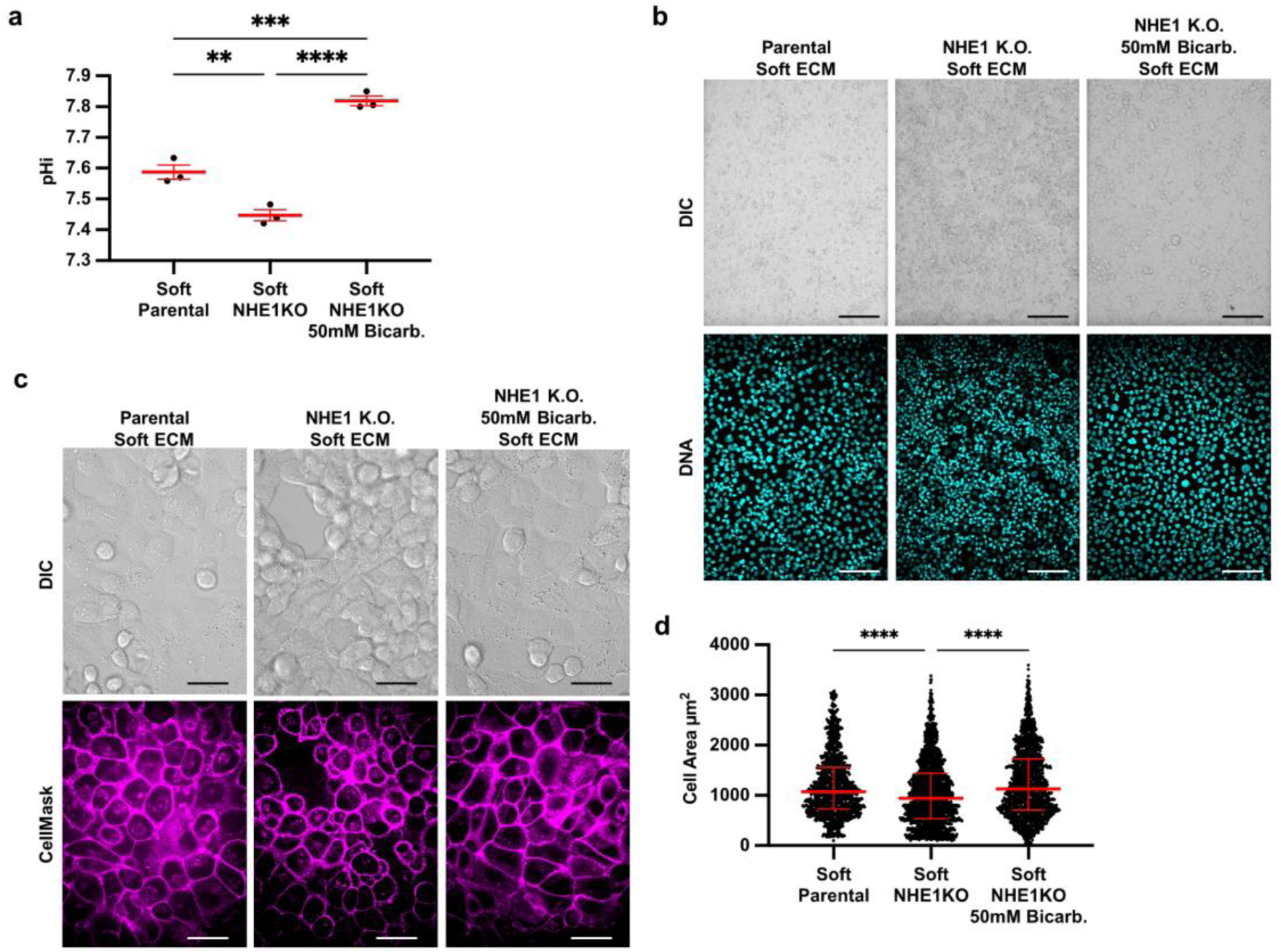
Low pHi is sufficient to induce vasculogenic mimicry on a soft ECM. **a)** Quantification of pHi data from parental H1299 cells and H1299 cells where NHE1 knockout via CRISPR (H1299-NHE1 K.O.). with and without treatment with sodium bicarbonate (Bicarb.) (see methods) (n=3 biological replicates. Red lines show means ± SEM). **b)** Representative images of H1299 cells plated on soft HA gels (0.5% PEGDA) with and without lowered pHi (H1299-NHE1 K.O.) and with or without increased pHi (H1299-NHE1 K.O. Bicarb.). Images show differential interference contrast (DIC) and Hoechst 33342 (DNA, cyan). Scale bars: 100 μm. **c)** Representative images of H1299 cells plated on soft HA gels (0.5% PEGDA) with and without lowered pHi (H1299-NHE1 K.O.) and with or without increased pHi (H1299-NHE1 K.O. Bicarb.). Images show differential interference contrast (DIC) and CellMask Deep Red membrane stain (Cy5, magenta). Scale bars: 50 μm. **d)** Quantification of single-cell area collected as shown in (a) (n=3 biological replicates, n=383 parental, n=267 NHE1 K.O., n=315 NHE1 K.O. Bicarb. Red lines show medians ± IQR).

## Discussion

Our work identifies pHi dynamics as a previously unrecognized regulator of stiffness-dependent VM in metastatic cancer cell models. We combined physiologically relevant tunable-stiffness hydrogel systems with single-cell pHi imaging and quantitative microscopy approaches to reveal novel molecular integration of the extracellular mechanical environment and pHi. We show that increasing ECM stiffness, driven by either increased protein concentration or crosslinking, lowers the single-cell pHi of lung, breast, and bone cancer cell lines. Most previously described tumorigenic behaviors such as hyperplasia^55^, metastatic progression^35,56^, and drug resistance^56^ are associated with increased pHi. However, recent work suggests a potential role for comparatively low cancer cell pHi in regulating hypoxia response^57^ and modulating tumor initiating cell phenotypes^14^. Adding to these recent data, our new findings show that low pHi in cancer cells is both a necessary and sufficient driver of VM. When we raise pHi in cells plated on stiff ECM, we attenuate VM phenotypes, overriding mechanosensitive regulation of VM. More surprisingly, low pHi was sufficient to drive formation of VM phenotypes on soft ECM (in the absence of mechanical stiffening).

Our work characterizing pH-dependent molecular drivers of VM identified β- catenin as a pH-sensitive regulator of vasculogenic mimicry. We also show that another VM regulator, FOXC2, has stiffness-dependent but pHi-*independent* activity. Our molecular characterization of VM regulators using tunable-stiffness hydrogels in combination with pHi manipulation approaches also resolved conflicting data in the literature on the interdependence of β-catenin and FOXC2 in VM regulation. Importantly, our data suggest that β-catenin is a necessary regulator of stiffness-dependent VM and that FOXC2 transcriptional activation is not sufficient to drive VM in the absence of stabilized β-catenin.

Our data reveal that pHi is a master regulator of VM and can override mechanosensitive phenotypes in 2D, improving understanding of molecular mechanisms driving cancer cell adaptive behaviors in the context of a stiffening ECM. In this work, we limit our characterization to 2D ECM models and a handful of metastatic cell models while focusing on just one tumorigenic mechanosensitive phenotype (VM). This approach enables us to combine quantitative single-cell pHi measurements and pHi manipulation with tunable-stiffness hydrogel systems to differentiate the contributions of stiffness and pHi in these complex cell morphology phenotypes. Our findings provide the groundwork for future experiments investigating pHi-dependent mechanosensitive behaviors in more complex 3D tumor spheroid models or even co-culture models with cancer-associated fibroblasts or immune cells. Prior work has already independently shown that complex 3D environments produce increased vasculogenic mimicry and phenotypic heterogeneity^58^ and pronounced pHi gradients^59^. Our current findings motivate expanding these studies to more complex mechanical and cellular environments to explore mechanistic roles for pHi dynamics in regulating other mechanosensitive tumorigenic behaviors such as durotaxis, invasion, and phenotypic plasticity.

## Methods

### Cell Culture

H1299 (parental ATCC CRL-5803) or H1299-NHE1 K.O. (CRISPRed cell line was a gift from Dr. Diane Barber at the University of California, San Francisco) were grown in RPMI 1640 (Corning, 10-040-CV) supplemented with 10% Fetal Bovine Serum (FBS, Peak Serum, PS-FB2). When using CHIR treatment (CHIR 99021, Ambeed, A133052), 10uM was supplemented to media 24 hours after plating.

MDA-MB-231 (ATCC HTB-26) cells were grown in DMEM (Corning, MT10013CVV) supplemented with 10% FBS. All cells were maintained at 5% CO_2_ and 37°C in a humidified incubator. To increase pHi, cells were cultured under normal conditions for 24 hours before being treated for 24 hours with culture media supplemented to a final concentration of 50-100mM Sodium Bicarbonate (Sigma-Aldrich; S6297-250G).

U2 OS (ATCC HTB-96) cells were grown in McCoy’s 5A (Modified) Medium (ThermoFisher Scientific; 16600082) supplemented with 10% FBS. All cells were maintained at 5% CO_2_ and 37°C in a humidified incubator.

### Transient Expression and Stable Cell Line Generation

H1299 and MDA-MB-231 mCherry-pHluorin expressing cells were generated as previously described^35^. FOXC2-TAG-Puro (LipExoGen Biotech, SKU:LTV-0061) positive H1299 cells were generated using lentiviral article transduction. Briefly, H1299 cells were plated at 50% confluency in a 6 well tissue culture treated plate. After 24 hours, media was replaced with fresh media containing 10ug/mL of polybrene and 50uL/well of FOXC2-TAG-Puro lentiviral particles. Cells were incubated for 72 hours prior to selection with 0.8 mg/mL blasticidin (Thermo Fisher Scientific, BP264725). After 4 weeks of selection, GFP positive cells were sorted on a BD FACS ARIA III cell sorter using 488nm excitation with 515nm-545nm emission filter. These cells were collected into 1mL 1XPBS using high purity sort settings. Cells were then centrifuged and plated in complete RPMI media with 0.8 mg/mL blasticidin.

### Preparation of tunable-stiffness hydrogels

#### Matrigel or Geltrex gel systems

Matrigel (Corning 356231, Lot 9035003) or Geltrex (Gibco, A14132-02, add LOT) coated plates were made in 35 mm diameter, 4-well (9.5 mm/well) glass bottom dishes (Matsunami, D141400). Stock Matrigel or Geltrex (12 or 16 mg/mL respectively) were diluted in cold complete media to concentrations of 4 mg/mL, 6 mg/mL, and 8 mg/mL which cover a range of stiffness from 50 Pa to ∼1000 Pa^30–32^. Each well was coated with 2.6 µL matrix per mm of well surface area (25 µL/well for 9.5 mm 4-well plate). Matrix was allowed to solidify at 37°C for 20 minutes prior to cell plating. Cells were plated at 5,000 cells per well in 100 µL solution volume.

#### HA gel system

HyStem-C (Advanced BioMatrix GS313) gels are composed of thiol-modified hyaluronic acid (Glycosil, GS222F), thiol-modified gelatin (Gelin-S, GS231F), polyethylene glycol diacrylate (PEGDA, Extralink, GS3007F), and degassed, deionized water (DG Water)^22,30^. Basement matrix solution was made of 1:1 Glycosil and Gelin-S and varying final PEGDA percentages (0.5, 1, 2, and 4%) were prepared in degassed, deionized water. The basement matrix solution and respective percentage PEGDA were mixed in a 4:1 parts ratio immediately before plating. Each well was coated with 1.4 µL matrix per mm of well surface area (13.5 µL/well). Cells were plated on the pre-prepared synthetic ECM plates 48 hours prior to imaging at 5,000 (single-cell pHi measurements) or 75,000 (VM imaging/staining) cells/well in 100 µL solution volume. HA gels were pre-prepared a maximum of 3 days prior to plating of cells, and stored with Dulbecco’s phosphate buffered saline (DPBS) (Quality Biological, 114-057-101) in each well to maintain hydration at 4°C.

#### Microscope System

Confocal images were collected on a Nikon Ti-2 spinning disk confocal with a 10x (PLAN APO NA0.45) air objective, 40x (CFI PLAN FLUOR NA1.3) oil immersion objective, and 60x (PLAN APO NA1.4) oil immersion objective. The microscope is equipped with a stage-top incubator (Tokai Hit), a Yokogawa spinning disk confocal head (CSU-X1), four laser lines (405 nm (100 mW), 488 nm (100 mW), 561 nm (100 mW), 640 nm (75 mW)), a Ti2-S-SE motorized stage, multi-point perfect focus system, and an Orca Flash 4.0 CMOS camera. Images were acquired under the following settings: pHluorin (GFP) (488 nm laser excitation, 525/36 nm emission), mCherry (561 nm laser excitation, 630/75 nm emission), Cy5 (647 nm laser excitation, 705/72 nm emission), Hoechst 33342 Dye (405 nm laser excitation, 455/50 nm emission), TxRed (561 nm laser excitation, 605/52 nm emission), and SNARF (561 nm laser excitation, 705/72 nm emission) and differential interference contrast (DIC) were used. Acquisition times for each fluorescence channel ranged from 50-600 milliseconds.

#### Single-cell pHi measurements using mCherry-pHluorin and confocal microscopy

Prior to imaging, stage top incubator and microscope objectives were pre heated to 37°C and kept at 5% CO_2_/95% air. Single-cell pHi measurements were performed as previously described^35^. Briefly, initial fields of view (FOV) were collected on the cells in their respective media. Two isotonic buffers (25 mM HEPES, 105 mM KCl, 1 mM MgCl_2_) were prepared and supplemented with 10 μM nigericin (Thermo Fisher Scientific, N1495). For standardization, isotonic buffers were pre-warmed to 37°C and pH of the “Nigericin buffers” was adjusted to ∼6.7 and ∼7.7 (with 1M KOH) (recorded for each biological replicate to the hundredths place). For each standardization point, cells were washed three times consecutively with no waiting time with appropriate Nigericin buffer followed by a 5-7 minute equilibration prior to image acquisition. All required buffer exchanges were carried out on the stage incubator to preserve XY positioning. Multiple Z-planes were collected with the center focal plane maintained using the Perfect Focus System (PFS).

#### Single-cell pHi measurements using SNARF-AM and confocal microscopy

U-2 OS cells were plated at 5x10^3^ cells per well (4-well glass bottom dish) in complete medium 48 hours prior to imaging. Cells were treated for pHi manipulation as described above. Cells were dye loaded in conditioned media with 10 μM of 5-(and-6)-Carboxy SNARF™-1 Acetoxymethyl Ester, Acetate (Fisher Scientific, cat: C1272) prepared as a 100 μM stock solution dissolved in a 10% DMSO in DPBS for 15 minutes at 37°C and 5% CO_2_. Dye-containing media was removed, and plates were washed three times for 5 min each time with pre-warmed complete growth medium containing appropriate treatment. FOVs were selected by viewing cells in differential interference contract (DIC). Images in SNARF (10% laser power, 400 ms), TxRed (30% laser power, 400 ms), and DIC (200 ms) channels for each FOV. SNARF dye was calibrated using three pre-warmed Nigericin buffers (as described above).

#### pHi Image Quantification

NIS Analysis Software was used to quantify pHi. All images were background subtracted using a region of interest (ROI) drawn on glass coverslip (determined by DIC). Individual ROIs were drawn for each cell in each condition (initial, high pH nigericin, and low pH nigericin). For each cell ROI, mean pixel intensities were quantified and pHluorin/mCherry or SNARF/TxRed ratios were calculated in Microsoft Excel. For each cell, the nigericin standard fluorescence intensity values were used to generate single-cell standard curves where single-cell pHi was back-calculated based on nigericin buffer pH values reported to the hundredths.

#### Proliferation Assay

H1299 cells were plated at 1,000 cells/well in a 24 well tissue-culture treated plate on pre-prepared matrix (65 µL/well) (see *Preparation of tunable-stiffness hydrogels*). After 24 and 48 hours of culture, cells were lifted via trypsinization (0.25%, Corning, 25- 0530Cl) for 20 minutes and counted by hemocytometer.

### Immunofluorescence Staining

#### Fixed Cell Staining

H1299 and MDA-MB-231 cells were plated at 75,000 cells/well in 100 µL solution volume on the pre-prepared synthetic ECM plates. After 48 hours, the media was removed and a 3.7% Formaldehyde (Alfa Aesar, 50-000) solution in DPBS was added to each well and allowed to fix at room temperature for 10 minutes. Cells were washed 3x2 minutes with DPBS before a permeabilization solution (0.1% Triton-X (Fisher Scientific, 9002-93-1) in DPBS) was added to each well for ten minutes at room temperature (RT). The Triton-X permeabilization solution was removed and cells were washed 3x2 minutes with DPBS at RT before a blocking solution (1% BSA (Fisher Scientific, BP1600-100) was added to cells for one hour at RT with rocking. The blocking solution was removed and cells were washed 3x2 minutes with DPBS before primary antibody solutions were added to each well and incubated with rocking at 4°C overnight. Primary antibodies were prepared in 1% BSA with 0.1% Triton-X at 1:50 dilutions. Primary antibodies used were: β-catenin mouse (BD Biosciences, BDB610154) Cyclin-D rabbit (Cell Signaling Technology, 55506S), and FOXC2 rabbit (Cell Signaling Technology, 12974S). The following day, primary antibody solutions were removed and cells were washed 3x2 minutes with DPBS before secondary antibodies (Goat anti-mouse IgG (H+L) Cross-Absorbed Secondary Antibody, Alexa Fluor 488; Invitrogen; A-11001, Goat anti-rabbit IgG (H+L) Secondary Antibody, Alexa Fluor 488; Invitrogen; A-11008) were added at 1:1,000 in solution of 1% BSA, 0.1% Triton-X, and Hoechst 33342 (DAPI; Thermo Scientific, cat: 62249 were added to each well (1:20,000) in DPBS and incubated with rocking at RT for one hour. Cells were washed 3x2 minutes with DPBS just prior to imaging on the Nikon Ti-2 spinning disk confocal with a 40x oil immersion objective. Images were captured with multiple Z planes to allow visualization of labeled protein colocalization. After acquisition, IMARIS Software (Bitplane, Oxford Instruments, version 9.5.1) and Nikon Elements Analysis software were used to quantify stained proteins. Nuclear pools of proteins were identified using IMARIS software by generating surfaces based on the DAPI channel that represent individual cell nuclei. Mean intensities for all channels within each nuclear surface were exported and analyzed for statistical significance using GraphPad Prism software. Whole cell protein abundance was determined by drawing regions of interest in Nikon Elements Analysis software of single cells. Mean intensities for all channels were exported and analyzed for statistical significance using GraphPad Prism software.

#### Live Cell Staining

H1299, MDA-MB-231, and U-2 OS cells were plated on the pre-prepared synthetic ECM plates 48 hours prior to imaging at 75,000 cells/well in 100 µL solution volume. Images were acquired as outlined in the above sections. Cell nuclei and cell membranes were visualized via Hoechst dye (DAPI; Thermo Scientific, cat: 62249; 1:10,000) and CellMask Deep Red (Thermo Fisher, C10046; 1:20,000), respectively, incubated for 15 minutes at 37⁰ C in complete media. Fields of view were selected by visualizing nuclei (DAPI) and images were collected in the DAPI (30% laser power, 600 ms), GFP (30% laser power, 600 ms), Cy5 (30% laser power, 600 ms), and DIC (32.6 DIA, 50 ms) channels. Individual cells were analyzed by IMARIS software by generating cells based on the CellMask channel that represents cell membranes. Cell areas were exported and analyzed for statistical significance using GraphPad Prism software.

#### Single-cell FOXC2 transcriptional activity assay using live-cell microscopy

FOXC2-TAG-Puro expressing H1299 cells were plated on the pre-prepared synthetic ECM plates 48 hours prior to imaging at 75,000 cells/well in 100 µL solution volume. Images were acquired as outlined in the above sections. Cell nuclei and cell membranes were visualized via Hoechst dye (DAPI; Thermo Scientific, cat: 62249; 1:10,000) and CellMask Deep Red (Thermo Fisher, C10046; 1:20,000), respectively, incubated for 15 minutes at 37⁰ C in complete media. Fields of view were selected by visualizing nuclei (DAPI) and images were collected in the DAPI (30% laser power, 600 ms), GFP (30% laser power, 600 ms), Cy5 (30% laser power, 600 ms), and DIC (32.6 DIA, 50 ms) channels. Whole-cell regions of interest (ROIs) were drawn within individual cells using cell mask as a membrane marker and the average GFP intensity for individual cells were exported to Excel. Single-cell intensities were imported to GraphPad Prism for statistical analysis and visualization.

#### BCECF plate reader assays

Cells were plated at 4.0×10^4^–8.0×10^4^ cells/well in a 24-well plate and incubated overnight. Cells were treated with 2 μM 2′,7′-bis-(2-carboxyethyl)-5-(and-6)- carboxyfluorescein, acetoxymethyl ester (BCECF-AM; VWR, 89139-244) for 20 min at 37°C and 5% CO_2_. H1299 parental and NHE1 K.O. cells were washed three times for 5 min each time with a pre-warmed (37°C) HEPES-based wash buffer (30 mM HEPES pH 7.4, 145 mM NaCl, 5 mM KCl, 10 mM glucose, 1 mM MgSO_4_, 1 mM KHPO_4_, 2 mM CaCl_2_, pH 7.4) to match their low bicarbonate medium (RPMI) and NHE1 K.O. Bicarb. cells were washed three times for 5 min each time with a pre-warmed (37°C) HEPES- based wash buffer (30 mM HEPES pH 7.4, 95 mM NaCl, 5 mM KCl, 10 mM glucose, 1 mM MgSO_4_, 1 mM KHPO_4_, 2 mM CaCl_2_, pH 7.4) to match sodium bicarbonate treatment. For standardization, three calibration buffers (25 mM HEPES, 105 mM KCl, 1 mM MgCl_2_) were supplemented with 10 μM nigericin (Thermo Fisher Scientific, N1495), pH was adjusted to ∼6.7, ∼7.0, and ∼7.7, and were pre-warmed to 37°C. Fluorescence was read (excitation of 440 and 490 nm, both with emission at 535 nm) on a Cytation 5 (BioTek) plate reader incubated at 37°C with 5% CO_2_. Kinetic reads were taken at 15-s intervals for 5 min, using a protocol established within BioTek Gen5 software. After the initial pHi read, the HEPES/bicarbonate wash was aspirated and replaced with one of the nigericin buffer standards, and cells were incubated at 37°C with 5% CO_2_ for 7 min. BCECF fluorescence was read by the plate reader as above. This process was repeated with the second nigericin standard. As it takes significant time to equilibrate CO_2_ in the plate reader, we did not measure nigericin standardizations without CO_2_. The mean intensity ratio (490/440 values) was derived from each read. Measurements were calculated from a nigericin linear regression using exact nigericin buffer pH to two decimal places^37^.

#### NHE1 Recovery Assay

80,000 cells were plated in the first two rows of a 24-well plate two days prior to transfection (one row of H1299 parental, the other H1299 NHE1 K.O.). Cells were treated with 2 μM 2′,7′-bis-(2-carboxyethyl)-5-(and-6)-carboxyfluorescein, acetoxymethyl ester (BCECF-AM; VWR, 89139-244) for 20 min at 37°C and 5% CO_2_. H1299 parental and NHE1 K.O. cells were washed three times for 5 min each time with a pre-warmed (37°C) HEPES-based wash buffer (30 mM HEPES pH 7.4, 145 mM NaCl, 5 mM KCl, 10 mM glucose, 1 mM MgSO_4_, 1 mM KHPO_4_, 2 mM CaCl_2_, pH 7.4) to match their low bicarbonate medium (RPMI). Fluorescence was read (excitation of 440 and 490 nm, both with emission at 535 nm) on a Cytation 5 (BioTek) plate reader incubated at 37°C with 5% CO_2_. Kinetic reads were taken at ∼30 sec intervals for 5 min, using a protocol established within BioTek Gen5 software. Initial baseline images were taken in the HEPES buffer at pH 7.4 (6 mins total). Next, cells were loaded with ammonium chloride using a HEPES-based ammonium chloride buffer (30mM HEPES pH-7.4, 30mM NH_4_Cl, 115mM NaCl, 5mM KCl, 10mM glucose, 1mM MgSO_4_, 1mM KHPO_4_, 2mM CaCl_2_) and cells were read for 6min. An acid load was induced by removing the ammonium chloride buffer and replacing it with the HEPES buffer (no NH_4_Cl). Cells were read while they recovered for 10 minutes. A calibration curve was then obtained by imaging the cells in nigericin containing buffers at the various pH values, around 7.5, 7.0, and 6.5. The standard curve was then used to back calculate the pHi of the cells during the experiment as previously described^35^. The data was normalized to the initial point of the recovery period to look at the recovery rate differences.

#### Statistical analysis

GraphPad Prism was used to prepare graphs and perform statistical analyses. All data sets were subject to normality tests (D’Angostino & Pearson, Anderson-Darling, Shapiro-Wilk, and Kolmogorov-Smirnov) and outlier analyses using the ROUT method (Q=1%). For non-normally distributed data, a Kruskal–Wallis test with Dunn’s multiple comparisons correction was used and variance is reported as IQR (Figures 1-6). For fold increase in cell number and population pHi data, one-way ANOVA was used and variance is reported as SEM (Figure 2E, 6A). One-way ANOVA tests assume equal variance while Kruskal- Wallis does not. All significance was indicated in figures by the following: \**P*<0.05; \*\**P*<0.01; \*\*\**P*<0.001; \*\*\*\**P*<0.0001.

#### Replication

We performed 3-6 biological replicates for all cell-based data. Because single-cell imaging is used heavily, we also confirmed pHi manipulation across each replicate included for subsequent VM or molecular marker analyses.

#### Blinding

In addition to the quantitative analysis of vascular mimicry, we blinded imaging data and had other members of the White Lab group/score based on morphology changes. In all cases, White lab members were able to reliably differentiate between soft and stiff matrix but were unable to reliably differentiate soft and “stiff with high pHi” samples.

#### Considering sex as a biological variable

This is not appropriate for this experimental data.

#### Antibodies

Where possible, antibodies were selected from those previously validated by the authors using genetic validation techniques (knock-out of protein of interest, over- expression of protein of interest, selection of cell line that doesn’t express protein as true negative control) and independent antibody validation techniques (comparison of staining patterns of two independent antibodies and comparison to literature).

#### Cell Line Authentication

All actively cultured cell lines are tested every 6 months for mycoplasma, and genetically validated through STR Profiling every 5 years. Upon receipt of cells that have not been cultured in the lab, cell lines are tested for mycoplasma and genetically validated through STR Profiling. The White Lab’s response to any positive mycoplasma test is to discard any stocks made after the last mycoplasma-free testing. These cells were validated in 2022 and again in Dec 2024. If testing reveals that any of the commercially obtained cell lines (or cell lines from other labs) are misidentified, stocks will be discarded, anyone with whom the stocks were shared will be notified, and source (ATCC or another lab) will be notified.

## Online Supplementary Materials

Online supplementary materials include Figure S1- S9.

## Data and Materials Availability

All data are available in the main text or the supplementary materials.

## Acknowledgements

We like to thank members of the White lab for their helpful conversations and feedback during figure and manuscript preparation. We also thank Dr. Diane Barber for her generous gift of the H1299-NHE1 K.O. cell line. The spinning disk confocal microscope used in this work is a part of the Notre Dame Integrated Imaging Facility (NDIIF).

## Author contributions

Conceptualization: LML, KAW, DH-P; Methodology: LML, KAW, LA, KJT, DH-P; Validation: LML, KAW, DH-P; Formal analysis:, LML, LA, LNM, EH, JH, RM, IA, KJT, KAW, DH-P; Investigation: LML, LA, LNM, EH, JH; Resources: KAW, DH-P.; Data collection: LML, LA, LNM, EH, JH, IA, KJT; Data curation: LML, LA, LNM, EH, JH, IA, KJT, KAW, DH-P; Writing – original draft: LML, KAW; Writing - review & editing: LML, LA, LNM, EH, JH, RM, IA, KJT, DH-P; Visualization: LML, LNM, KAW, DH-P; Supervision: KAW, DH-P; Project administration: KAW, DH-P; Funding acquisition: LML, KAW, DH-P.

