## Supplemental Figures for "Intracellular pH dynamics respond to extracellular matrix stiffening and mediate vasculogenic mimicry through β-catenin"

**This PDF file includes:**

Supplementary Figures S1 to S9

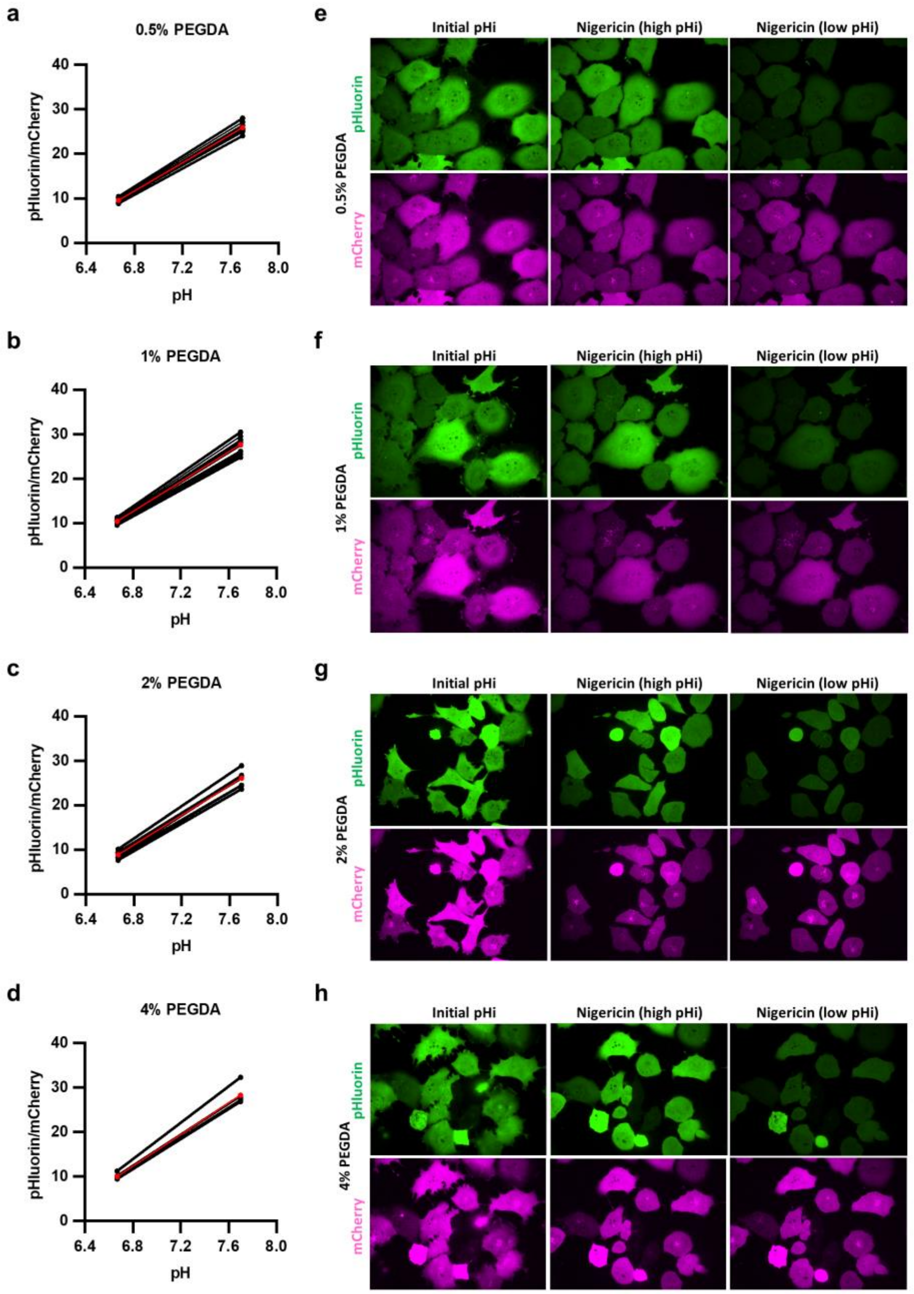


**Figure S1: Nigericin standardization of mCherry-pHluorin biosensor in H1299 cells.** Nigericin standard linear regressions across a single replicate for each single cell (black) and mean (red) for H1299-mCherry-pHluorin plated on **a)** 0.5% PEGDA (n=10), **b)** 1% PEGDA (n=10), **c)** 2% PEGDA (n=13), **d)** 4% PEGDA (n=7) HA gels. Single channel images of pHluorin (green) and mCherry (magenta) fluorescence at initial pHi imaging, high nigericin, and low nigericin standardizations for **e)** 0.5% PEGDA, **f)** 1% PEGDA, **g)** 2% PEGDA, **h)** 4% PEGDA. LUTs are identical across each cell line. Note that in all cases, single-cell nigericin standard curves were used to back-calculate single-cell pHi.

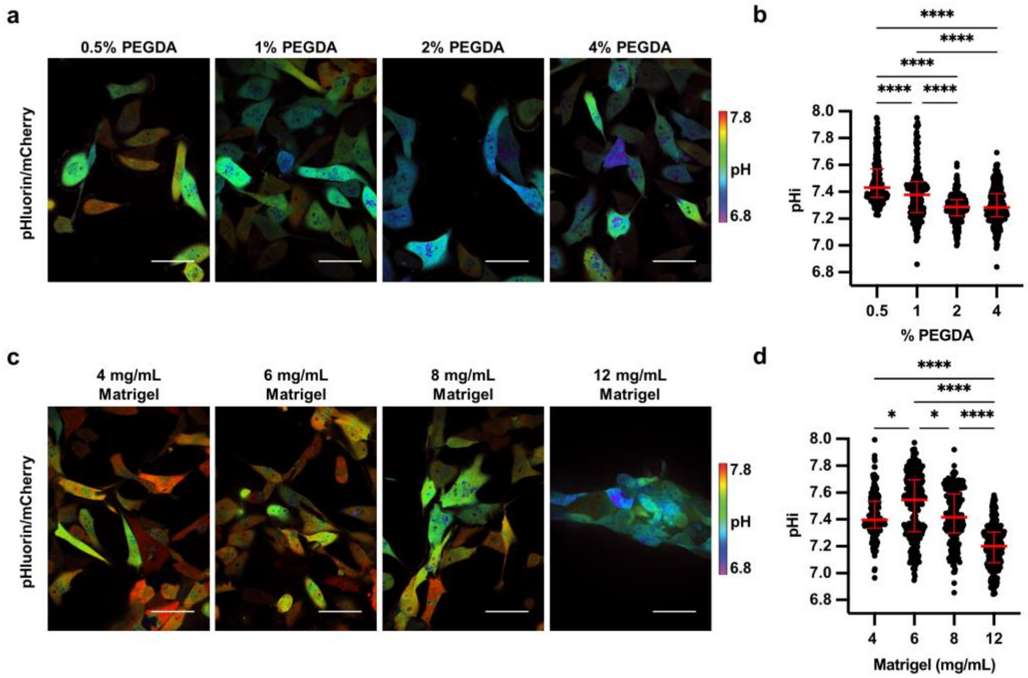


**Figure S2: Stiffening extracellular matrix lowers pHi in metastatic human breast carcinoma (MDA-MB-231). a)** Representative images of MDA-MB-231 cells stably expressing mCherry-pHluorin pH biosensor plated on varying HA gel stiffnesses. Images show ratiometric display of pHluorin/mCherry fluorescence. Scale bars: 50 μm. **b)** Quantification of single-cell pHi data collected as shown in (a) (n=3 biological replicates, n=194 0.5% PEGDA, n=228 1% PEGDA, n=265 2% PEGDA, n=334 4% PEGDA. Red lines show medians ± IQR). **c)** Representative images of MDA-MB-231 cells stably expressing mCherry-pHluorin pH biosensor plated on varying Matrigel stiffnesses. Images show ratiometric display of pHluorin/mCherry fluorescence. Scale bars: 50 μm. **d)** Quantification of single-cell pHi data collected as shown in (c) (n=3 biological replicates, n=210 4mg/mL, n=291 6mg/mL, n=222 8mg/mL, n=292 12mg/mL. Red lines show medians ± IQR). For (c) and (d), significance was determined by a Kruskal-Wallis test (**P*<0.05; ***P*<0.01; ****P*<0. 001; *****P*<0.0001).

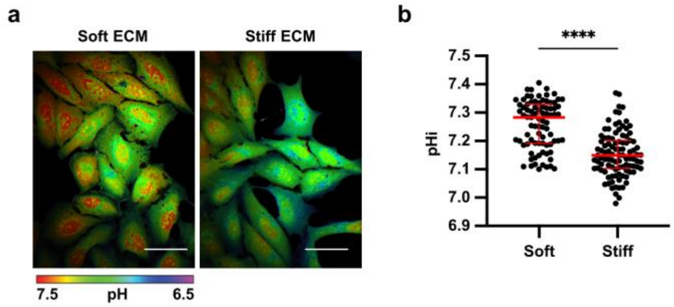


**Figure S3: Stiffening extracellular matrix increases pHi in metastatic bone cells (U-2 OS). a)** Representative images of U-2 OS cells loaded with SNARF pH dye plated on soft (0.5% PEGDA) and stiff (4% PEGDA) HA gels. Images show ratiometric display of SNARF/TxRed fluorescence ratios. Scale bars: 50 µm. **b)** Quantification of single-cell pHi data collected as shown in (a) (n=4 biological replicates; n=299 soft, n=305 stiff. Red lines show medians ± IQR). For (b), significance was determined by a Kruskal-Wallis test (*$P$<0.05; **$P$<0.01; ***$P$<0. 001; ****$P$<0.0001).

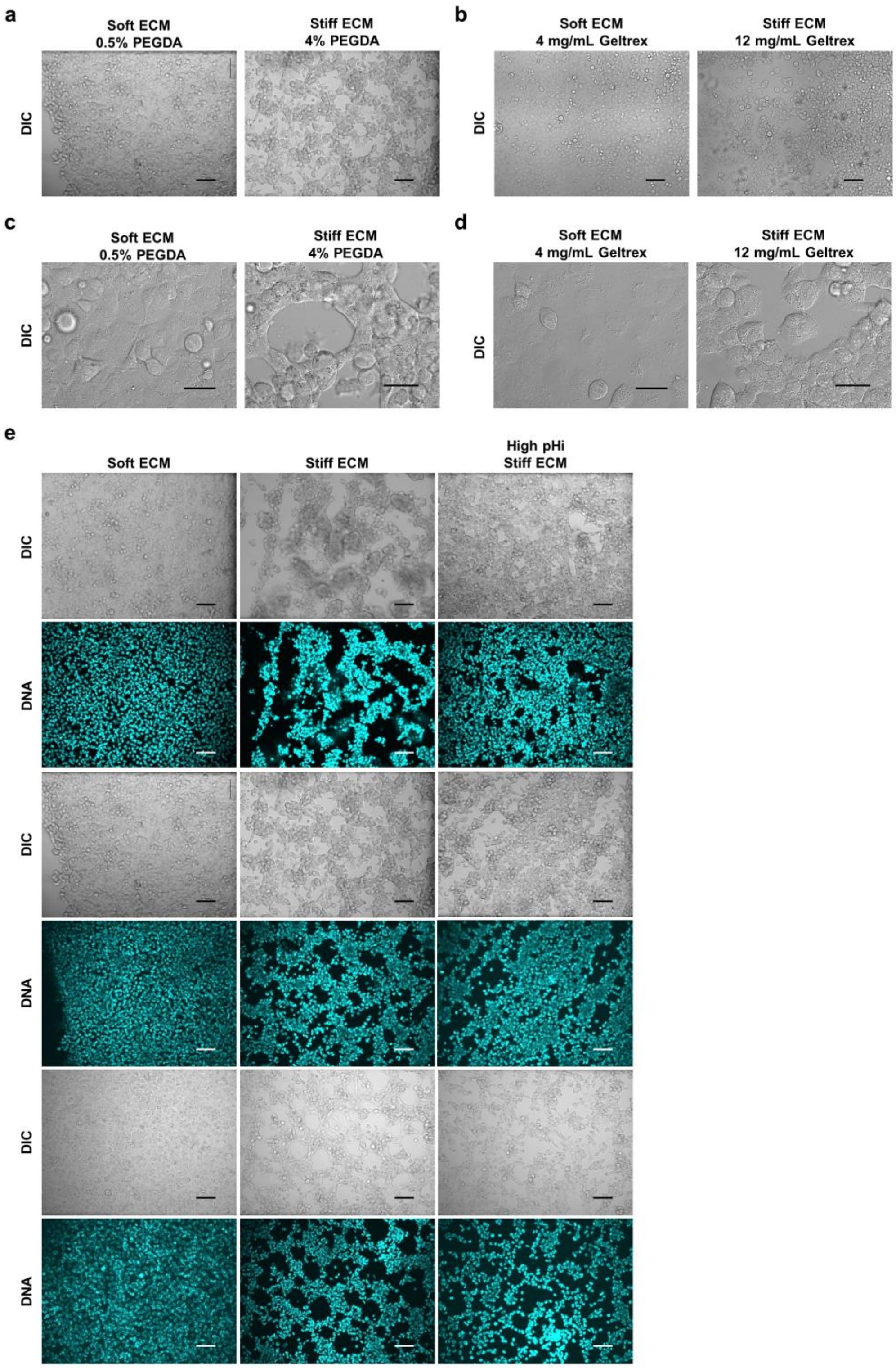


**Figure S4: Representative images of Vasculogenic mimicry phenotype on soft and stiff Geltrex and HA gels. a)** Differential interference contrast (DIC) images of H1299 cells plated on soft (0.5% PEGDA) and stiff (4% PEGDA) HA gels. Scale bars: 100 µm. **b)** Differential interference contrast (DIC) images of H1299 cells plated on soft

(4mg/mL) and stiff (12 mg/mL) Geltrex gels. Scale bars: 100 µm. **c)** Differential interference contrast (DIC) images of H1299 cells plated on soft (0.5% PEGDA) and stiff (4% PEGDA) HA gels. Scale bars: 50 µm. **b)** Differential interference contrast (DIC) images of H1299 cells plated on soft (4mg/mL) and stiff (12 mg/mL) Geltrex gels. Scale bars: 50 µm. **e)** Differential interference contrast (DIC) and Hoechst stain (DNA, cyan) images of H1299 cells plated on soft (0.5% PEGDA), stiff (4% PEGDA), and stiff (4% PEGDA) with high pHi HA gels. Scale bars: 100 µm.

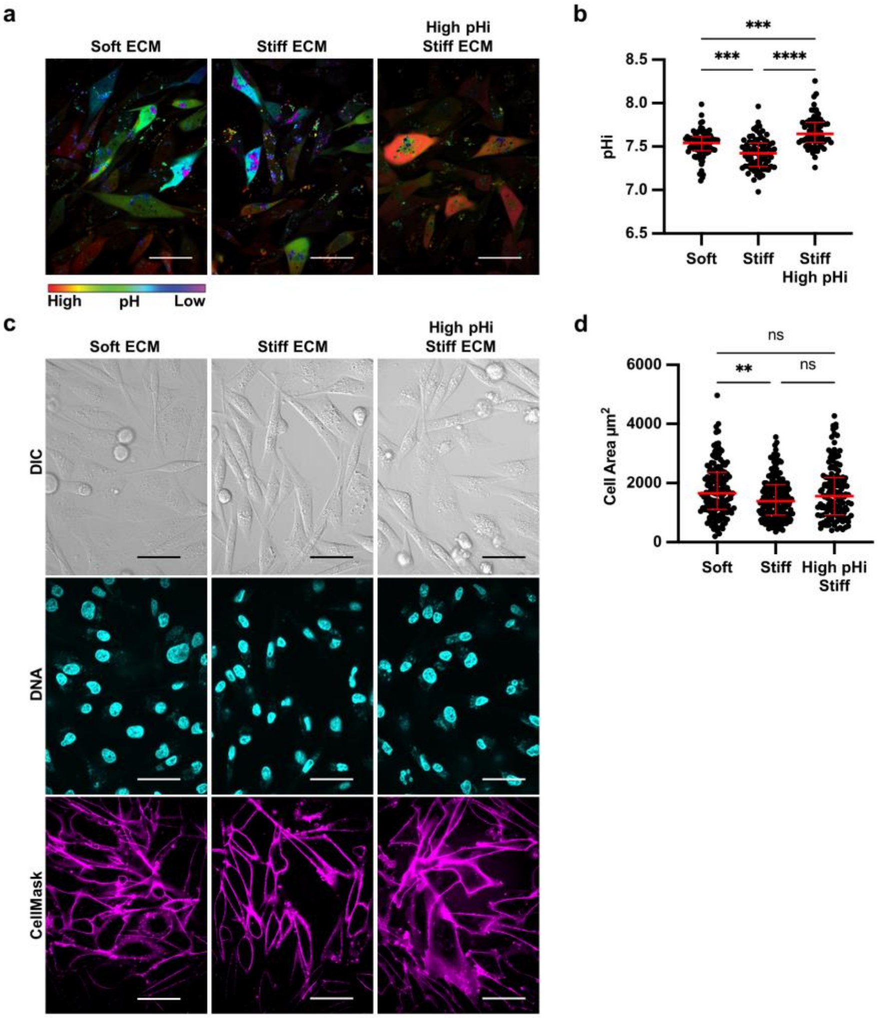


**Figure S5: Stiffness-dependent vasculogenic mimicry is reduced when pHi is increased on stiff ECM in metastatic breast cells (MDA-MB-231). a)** Representative images of MDA-MB-231 cells stably expressing mCherry-pHluorin pH biosensor plated on soft (0.5% PEGDA) and stiff (4% PEGDA) HA gels and stiff (4% PEGDA) with raised pHi (culture media supplementation with 50mM sodium bicarbonate). Images show ratiometric display of pHluorin/mCherry fluorescence ratios. Scale bars: 50 µm. **b)** Quantification of single-cell pHi data collected as shown in (a) (n=2 biological replicates; n=79 soft, n=81 stiff, n=69 stiff high pHi. Red lines show medians ± IQR). **b)** Representative images of MDA-MB-231 cells plated on soft (0.5% PEGDA) and stiff (4% PEGDA) HA gels and stiff (4% PEGDA) with raised pHi. Images show differential interference contrast (DIC), Hoechst 33342 (DNA, cyan) and CellMask Deep Red membrane stain (Cy5, magenta). Scale bars: 50 µm. **c)** Quantification of single-cell area collected as shown in (b) (n=2 biological replicates, n=151 soft, n=175 stiff, n=132 stiff high pHi. Red lines show medians ± IQR). For (b) and (d), significance was determined by a Kruskal-Wallis test (*$P$<0.05; **$P$<0.01; ***$P$<0. 001; ****$P$<0.0001).

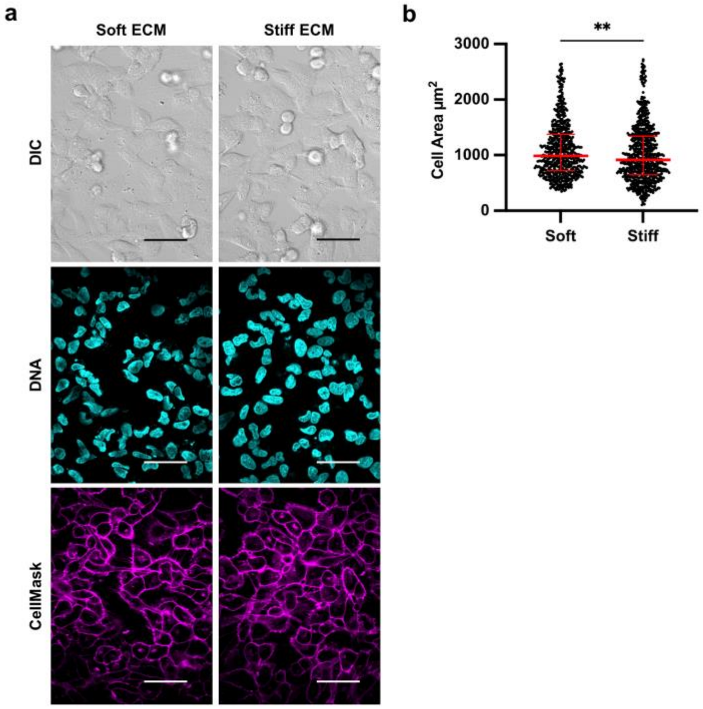


**Figure S6: Stiffness-dependent vasculogenic mimicry is reduced when pHi is increased on stiff ECM in metastatic bone cells (U-2 OS). a)** Representative images of U-2 OS cells plated on soft (0.5% PEGDA) and stiff (4% PEGDA) HA gels. Images show differential interference contrast (DIC), Hoechst 33342 (DNA, cyan) and CellMask Deep Red membrane stain (Cy5, magenta). Scale bars: 50 µm. **c)** Quantification of single-cell area collected as shown in (a) (n=2 biological replicates, n=855 soft, n=836 stiff. Red lines show medians ± IQR). For (b), significance was determined by a Kruskal-Wallis test (**P*<0.05; ***P*<0.01; ****P*<0. 001; *****P*<0.0001).

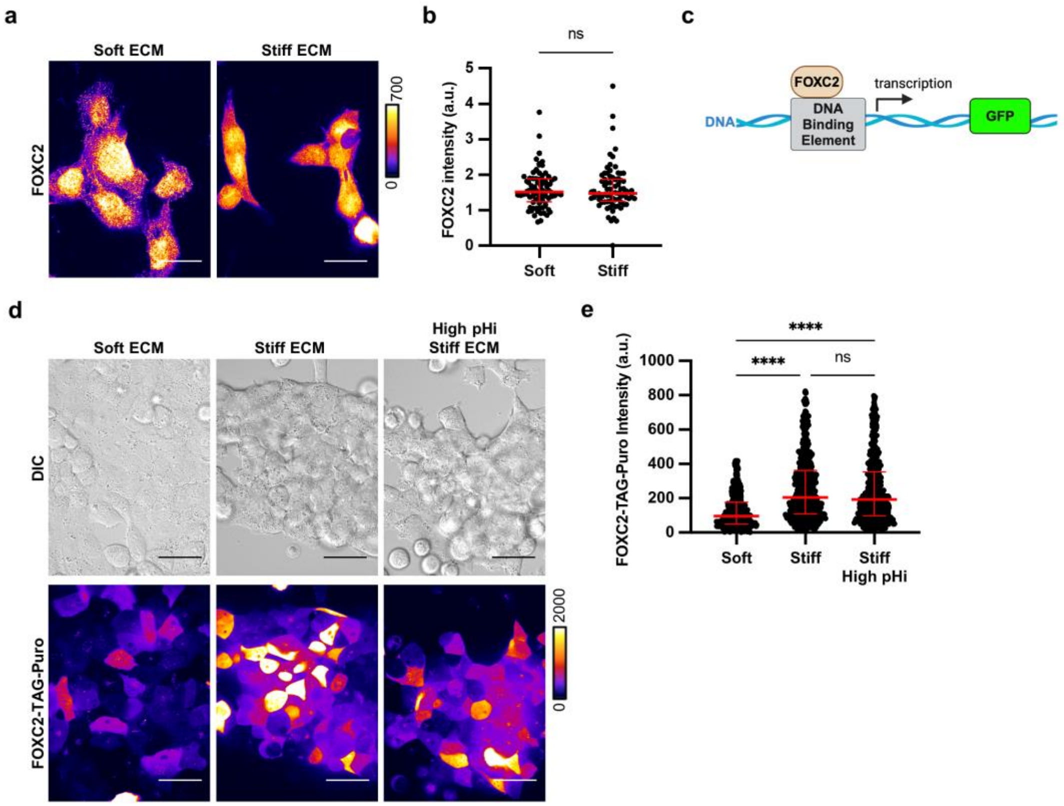


**Figure S7: FOXC2 has stiffness-dependent activity that is insensitive to pHi. a)**. Representative images of H1299 cells plated on soft (4 mg/mL) and stiff (12 mg/mL) Geltrex fixed and stained for FOXC2. FOXC2 is pseudocolored according to scale. Scale bars: 50 µm. **b)** Quantification of FOXC2 intensity per cell collected as shown in (a) (n=3 biological replicates, n=80 soft and n=77 stiff. Red lines show medians ± IQR). **c)** Schematic of FOXC2-TAG-Puro reporter of FOXC2 transcriptional activity. **d)** Representative images of H1299 cells plated on soft (0.5% PEGDA), stiff (4% PEGDA) and stiff (4% PEGDA) with raised pHi HA gels. Images show Brightfield display (DIC) and FOXC2-TAG-Puro. FOXC2-TAG-Puro is pseudocolored according to scale. Scale bars: 50 µm. **e)** Quantification of FOXC2-TAG-Puro intensity per cell collected as shown in (d) (n=3 biological replicates, n=416 soft, n=478 stiff, n=461 stiff high pHi. Red lines show medians ± IQR). For (b) and (e), significance was determined by a Kruskal-Wallis test (**P*<0.05; ***P*<0.01; ****P*<0. 001; *****P*<0.0001).

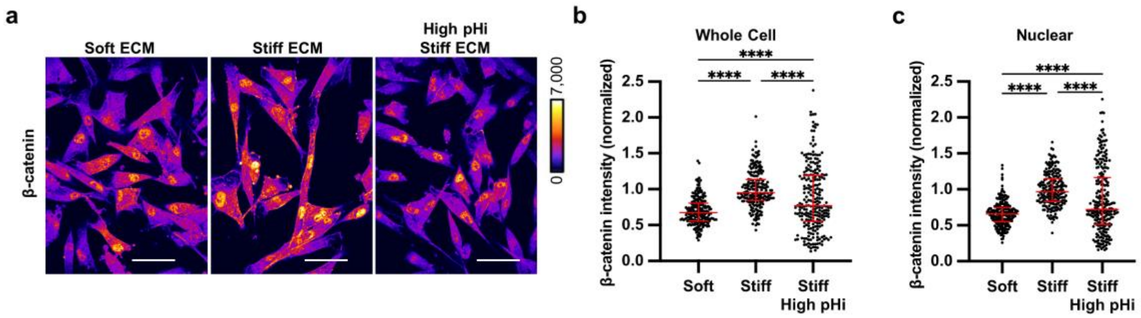


**Figure S8: Increased pHi reduced β-catenin abundance and nuclear localization in stiff matrix conditions. a)** Representative images of MDA-MB-231 cells plated on soft (0.5% PEGDA), stiff (4% PEGDA) and stiff with raised pHi (4% PEGDA) HA gels fixed and stained for β-catenin. β-catenin is pseudocolored according to scale. Scale bars: 50 µm. **b)** Quantification of whole cell β-catenin intensity collected as shown in (a). (n=3 biological replicates, n=208 soft, n=205 stiff, n=245 stiff high pHi. Red lines show medians ± IQR). **c)** Quantification of nuclear β-catenin intensity collected as described in (a). (n=3 biological replicates, n=208 soft, n=204 stiff, n=244stiff high. Red lines show medians ± IQR). For (b) and (c), significance was determined by a Kruskal-Wallis test (**P*<0.05; ***P*<0.01; ****P*<0. 001; *****P*<0.0001).

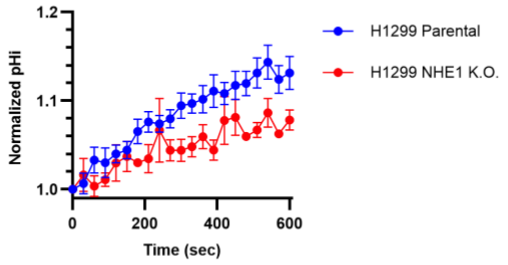


**Figure S9: NHE1 activity is decreased in H1299 cells with CRISPR knockout of NHE1.** Normalized pHi recovery measurements after acid load of parental H1299 cells (see methods) and H1299 cells where NHE1 has been removed via CRISPR (H1299-NHE1 K.O., see methods). (n=4 biological replicates; n=21 H1299 parental, n=21 H1299 NHE1 K.O.); (means ± SEM).
